# Coupling between ribotypic and phenotypic traits of protists across life-cycle stages and temperatures

**DOI:** 10.1101/2020.12.28.424523

**Authors:** Songbao Zou, Rao Fu, Huiwen Deng, Qianqian Zhang, Eleni Gentekaki, Jun Gong

## Abstract

Relationships between ribotypic and phenotypic traits of protists across life-cycle stages remain largely unknown. Herein, we used single cells of two soil and two marine ciliate species to examine phenotypic and ribotypic traits and their relationships across lag, log, plateau, cystic stages and temperatures. We found that *Colpoda inflata* and *C. steinii* demonstrated allometric relationships between 18S rDNA copy number per cell (CNPC), cell volume (CV), and macronuclear volume across all life-cycle stages. Integrating previously reported data of *Euplotes vannus* and *Strombidium sulcatum* indicated taxon-dependent rDNA CNPC–CV functions. Ciliate and prokaryote data analysis revealed that the rRNA CNPC followed a unified power-law function, only if the rRNA-deficient resting cysts were not considered. Hence, a theoretical framework was proposed to estimate the relative quantity of resting cysts in the protistan populations with total cellular rDNA and rRNA copy numbers. Using rDNA CNPC was a better predictor of growth rate at a given temperature than rRNA CNPC and CV, suggesting replication of redundant rDNA operons as a key factor that slows cell division. Single-cell high throughput sequencing and analysis after correcting sequencing errors revealed multiple rDNA and rRNA variants per cell. Both encystment and temperature affected the number of rDNA and rRNA variants in several cases. The divergence of rDNA and rRNA sequence in a single cell ranged from 1% to 10%, depending on species. These findings have important implications for inferring cell-based biological traits (e.g., species richness, abundance and biomass, activity, and community structure) of protists using molecular approaches.

**Importance:** Based on phenotypic traits, traditional surveys usually characterize organismal richness, abundance, biomass, and growth potential to describe diversity, organization and function of protistan populations and communities. The ribosomal RNA gene (rDNA) and its transcripts have been widely used as molecular markers in ecological studies of protists. Nevertheless, the manner in which these molecules relate to cellular (organismal) and physiological traits remains poorly understood, which could lead to misinterpretations of protistan diversity and ecology. The current research highlights the dynamic nature of cellular rDNA and rRNA contents, which tightly couple with multiple phenotypic traits in ciliated protists. We demonstrate that quantity of resting cysts and maximum growth rate of a population can be theoretically estimated using ribotypic trait-based models. The intra-individual sequence polymorphisms of rDNA and rRNA can be influenced by encystment and temperature, which should be considered when interpreting species-level diversity and community structure of microbial eukaryotes.

## Introduction

Protists are single-celled eukaryotes of astounding morphological and functional diversity inhabiting virtually every life-harboring niche on this planet. These microorganisms have complex life cycles comprising stages of distinct phenotypic traits (e.g., morphology, cell size, cystic/vegetative status, and growth rate), which undergo changes in response to fluctuating environmental variables (1-6). Recently, the advent of sequencing technologies has greatly facilitated the study of ribotypic traits, i.e. the qualitative and quantitative attributes of ribosomal (r)RNA genes (rDNA) and/or rRNA transcripts, which are used to assess protistan diversity and community composition in natural ecosystems (7).

The rDNA copy number (CN) and its dynamics provide important information on the diversity and quantity of protistan cells in molecular ecological studies. The eukaryotic rDNA comprises encoding regions for 18S, 5.8S, and 28S rRNA, non-coding internal transcribed regions (ITS1 and ITS2) and intergenic spacers and is generally arranged in tandem arrays of multiple copies. The transcribed 18S rRNA is the structural RNA for the small subunit of eukaryotic cytoplasmic ribosomes, which are the sites of protein synthesis and thus closely tied to cellular growth and development (8). Recently, variations in per-cell rDNA and rRNA CNs have been linked to phenotypic traits, such as cell volume (CV), and growth rate of exponentially growing ciliated protists (9). The dynamic nature of rDNA CN across life cycle stages has also been demonstrated in a foraminiferan species (10). Nevertheless, the association(s) between ribotypic and phenotypic traits in protists remains little explored. Moreover, the quantitative relations, if any, between ribotypic and phenotypic traits across different growth stages (e.g., lag, log, plateau and cystic phases) have yet to be investigated.

Cysts are dormant forms comprising essential life cycle stages for many protistan taxa exposed to unfavorable conditions (1, 11, 12). However, rDNA CN data of cysts are only available for a limited number of protistan species (e.g., 13, 14), and CN of rRNA in a cyst has seldomly been quantified, probably due to the common assumption that rDNA is sparsely or not at all transcribed in the dormant stages. The rRNA:rDNA CNs ratio has been used as an indicator of microbial activity/dormancy, however this approach is not entirely problem-free (9, 15). Experimental data of rRNA and rDNA CNs for different types of cysts are lacking. These can be used to build suitable models in order to properly estimate dormancy in environmental samples.

The rDNA operons of diverse protistan groups have a high degree of intra-individual sequence polymorphisms (16-21). During the life cycle of a protistan cell (10), and in response to changing temperatures (9), cellular rDNA copies are gained or lost. However, little is known about whether specific rDNA sequence variants are selected at different life cycle stages or under varying conditions. Similarly, the dynamics of rRNA sequence variants under these conditions warrant further investigation, as selective transcription of specific rDNA operons has been previously shown for several prokaryotes and a parasitic protist (e.g., 22-25).

Ciliates comprise one of the most intensely studied groups of protists making them ideal models for examining links between ribotypic and phenotypic traits across life history stages. Ciliate species have two morphologically and functionally distinct nuclei, a polyploid somatic macronucleus and one or several diploid germline micronuclei. The macronucleus contains hundreds to thousands of transcriptionally active nanochromosomes that function in all cellular events. The micronucleus is transcriptionally silent during cell growth, but following sexual reproduction it gives rise to the macronucleus (26). The latter contains numerous nucleoli, the sites of rDNA sequences, rRNA synthesis, and ribosome subunit assembly (27). During encystment and excystment, the number and size of nucleoli varies, accompanying loss and gain of macronuclear DNA (28, 29). This raises the question of whether in ciliates cellular ribotypic CNs are more related to macronuclear size than to CV across life cycle stages.

*Colpoda* has long been used as a model for the study of cyst-related biology and ecology (e.g., 28, 30, 31). In this work, we used two ubiquitously distributed soil ciliate species of *Colpoda* to investigate both rDNA and rRNA CNs across lag, log (exponential), plateau (stationary), unstable and resting cyst stages. The rDNA copies quantified and sequenced in this study were considered to be of macronuclear origin. Their phenotypic features (cell size, macronuclear size, nucleocytoplasmic ratio and maximum growth rate) were determined for cells reared at two temperatures. Temperature as a key environmental factor was investigated because of its importance in many aspects including climate change, effect on biochemical kinetics, metabolic and physiological processes, and modulation of body size and growth rate (32). The temperature and growth-stage relationships to intra-individual polymorphisms of ribotypic sequences were explored. We also extended previous work on the marine ciliates *Euplotes vannus* and *Strombidium sulcatum*, which were reared at three different temperatures by analyzing rDNA and rRNA sequence diversity in actively growing cells (9). Our main expectations were that: (1) the scaling relationships between rDNA and rRNA transcript CNs and phenotypic traits (CV and macronuclear size) would generally hold across life cycle stages including resting cysts; (2) the variant richness and composition of intra-individual rDNA and rRNA molecules would be life stage- and temperature-dependent.

## Materials and methods

### Source organisms, cultivation and treatments

Two cyst-forming, soil ciliate species, *Colpoda steinii* and *C. inflata*, were collected from a coastal apple orchard in Yantai, Shandong, China. The protists were cultivated and isolated using the non-flooded Petri dish method (33). *Euplotes vannus* and *Strombidium sulcatum* were originally isolated from the coastal waters off Qingdao (Shandong, China), and have been maintained in the laboratory since then. Cultivation of these ciliates and monitoring of their growth rate were as previously described (9). The *Colpoda* species were incubated for ten days at 18°C and 28°C and underwent distinct phases (lag, log, and plateau) during that time. Population growth at temperatures lower than 18°C or higher than 28°C was strongly inhibited, thus not investigated further. Following the plateau phase, both species consistently formed resting cysts, probably due to overpopulation and food depletion. Phase-specific cells (including the resting cysts) were individually isolated to determine both phenotypic and ribotypic traits at the single-cell level. Two marine species *E. vannus* and *S. sulcatum* were reared at 16°C, 21°C and 25°C. For these two taxa, only cells in log phase were investigated. In order to explore possible footprint effect of the higher temperature on ribotypic traits, an additional “touchdown” treatment (designated with “*”) was set as previously described (9). Briefly, a few individuals of the treatments maintained at 25 °C for 30 d were transferred and re-cultured at 16 °C for a month. All treatments were done in triplicate.

Many ciliate species form unstable cysts when temperature shifts beyond certain limitations in a short time period (30). These are generally larger than resting cysts, undergo complete ciliary dedifferentiation and return to vegetative cell state in a few hours. We tried obtaining unstable cysts of *Colpoda* by chilling cells for various periods of time. Tubes containing actively growing vegetative cells were transferred in three separate conditions: ice-water mixture (0°C) and chemostat reactors set at 4°C and 10°C. Twenty μL of culture solution were used to identify and count vegetative cells and unstable cysts using a stereomicroscope (XLT-400, Guiguang, China) at 45× magnification. Cultures were monitored every 0.5 to 2 h to determine the optimal time period for achieving the highest yield of unstable cysts. All chilling treatments were performed in triplicate. Since unstable cysts have high phenotypic plasticity, relevant phenotypic and ribotypic traits of this temporary stage were also determined and analyzed.

### Determination of growth rate, cell volume and macronuclear size

Cell abundances were determined once a day (Fig. 1A). Estimation of cell volume (CV) and calculations of growth rates of the two *Colpoda* species during the log phase were as previously described (9). At least 18 cells from each growth stage were randomly selected and fixed with Lugol’s solution (final concentration 2%). Fixed vegetative cells, resting and unstable cysts of both *Colpoda* species were ellipsoid, thus their cell volume was calculated according to the standard formula for a spheroid. Cell and macronuclear sizes of the *Colpoda* species were also determined based on specimens fixed with Bouin’s solution and stained using protargol impregnation (34). This approach reveals many morphological features of ciliates in detail, which enabled examination of the relationship between cell and macronuclear size.

**Figure 1.**
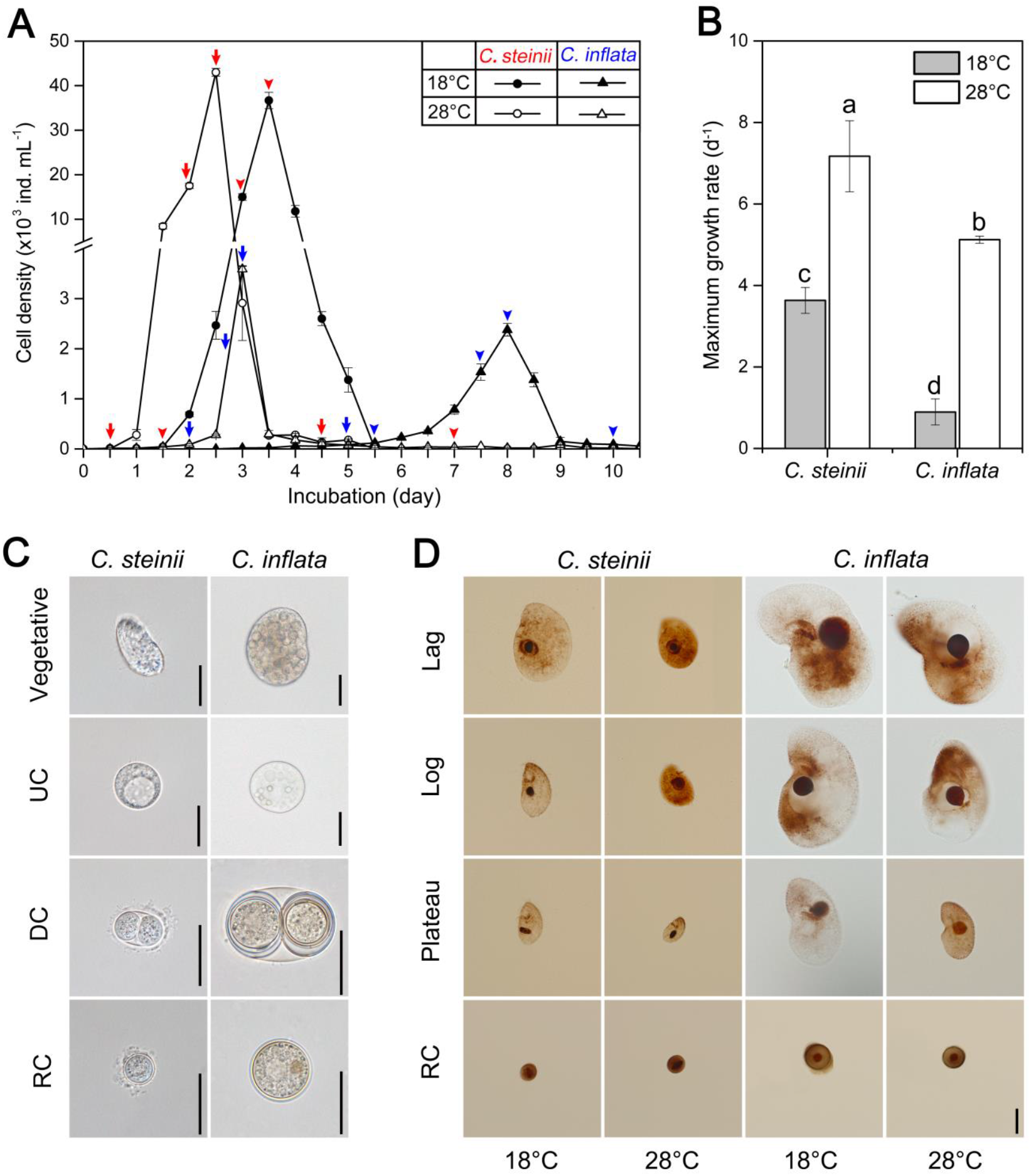
Growth and morphology of the soil ciliates *Colpoda steinii* and *C. inflata* reared at 18°C and 28°C. **(A)** Growth curves of *C. steinii* (arrows and arrowheads in red) and *C. inflata* (arrows and arrowheads in blue), depicting cell isolation time-points at lag, log, plateau and resting cyst phases. **(B)** Maximum growth rates of cells at log phase. Treatments not sharing common letters indicate significant differences (*P* < 0.05). **(C)** Morphology of vegetative cells, unstable cysts (UC), dividing cysts (DC) and resting cysts (RC) from life. **(D)** Microphotographs of fixed specimens after protargol impregnation. Cell and macronuclear sizes across life-cycle stages, temperatures and species. Scale bars = 20 μm.

### Cell lysis, DNA extraction and cDNA synthesis

Cell lysis, DNA extraction and RNA transcription of single-cells followed previous works (9). In brief, ciliate cells/cysts were washed and transferred into nuclease-free PCR tubes for cell lysis. Eighteen tubes (replicates) each containing a single intact cell were set up for cell lysis and nucleic acid extraction. Each cell was suspended in 1 μL of distilled water, followed by addition of 2 μL of mixed solution containing 1.9 μL 0.2% Triton-X-100 (Solarbio, Beijing, China) and 0.1 μL of recombinant RNase inhibitor (Takara, Biomedicals, Japan). The tubes were gently flicked, briefly centrifuged, and incubated at room temperature for 10 min. Nuclease-free PCR tubes, microtubes and micropippette tips (Axygen Scientific, CA, USA) were used for all manipulations. One μL of the cell suspension was used to extract genomic DNA, while 2 μL were used for RNA reverse transcription. Genomic DNA extraction was performed with a REDExtract-N-Amp Tissue PCR Kit (Sigma, St. Louis, MO), according to the manufacturer’s instructions, except the suggested volume of each reagent was modified to 1/50, as previously described (35).

The RNA reverse transcription of single cells was as previously described (9). The remaining 2 μL of cell suspension were mixed with 1 μL random hexamers B0043 (10 μM, Sangon, China) and 1 μL of dNTP mix (10 μM). Tubes were incubated at 72°C for 3 min and then transferred immediately onto ice. SuperScript III Reverse Transcriptase Kit (Invitrogen, CA, USA) was used for reverse transcription of RNA. The total volume used for PCR was 12.42 μL, which was made up of 4 μL rRNA template, 3 μL 5× first strand buffer, 0.3 μL SuperScript III reverse transcriptase, 1 μL 0.1M DTT, 2 μL Betaine (Sigma–Aldrich), 0.12 μL 0.5M MgCl_2_, 1 μL RNase inhibitor (Roche, Germany), and 1 μL DEPC-treated water. The following program was run on a thermal cycler (Biometra, Gottingen, Germany): an initial temperature of 50°C for 90 min, followed by 10 cycles at 55°C for 2 min and a final step at 70°C for 15 min. Both the DNA and synthesized cDNA were stored at -80°C for subsequent assays.

### Quantitative real-time PCR (qPCR)

To quantify 18S rDNA and rRNA (cDNA) copy numbers in *Colpoda* cells, the newly designed species-specific primers CsQ207f (5’-TAACCCTGGCAACAGGA-3’) and CsQ459r (5’-TGCAATCTCGCAACCCCA-3’) were used in qPCR assays to amplify a 252-bp fragment of the 18S rRNA gene of *C. steinii*. The eukaryote-specific primers EUK345f (5’-AAGGAAGGCAGCAGGCG-3’) and EUK499r (5’-CACCAGACTTGCCCTCYAAT-3’) (36) were used to amplify a 149-bp fragment of *C. inflata*. Two standard curves were constructed with the sequences of the two *Colpoda* species as previously described (19). All the qPCR reactions were performed on a 7500 Fast Real-Time PCR System (Applied Biosystems) with the following program: 95°C for 7 min, followed by 45 cycles of 95°C for 30 s, 55°C (60°C for *C. inflata*) for 1 min, and 72°C for 1 min (77°C for 25 s for *C. inflata*). The data were collected at 72°C and 77°C for *C. steinii* and *C. inflata*, respectively; all reactions ended with a melt curve stage from 60°C to 95°C (gradually increasing by 0.3°C). Standard curves were generated using log _10_ number of copies vs. the threshold cycle (Ct). The goodness of fit (*R*^2^) ranged from 0.998 to 1.000, with the amplification efficiency ranging from 98% to 108.9% (Fig. S1). All qPCR reactions were performed in triplicate. Controls without templates resulted in undetectable values for all samples. As the samples of rRNA also contained genomic rDNA, the rRNA (cDNA) copy numbers were calculated by subtracting the rDNA CNs from the sum of rDNA and cDNA copy numbers.

### Single-cell high throughput sequencing

The obtained genomic DNA and synthesized cDNA from single cells of each of the four ciliate species was used for high throughput sequencing. These included cells of *Colpoda steinii* and *C. inflata* at two typical stages (i.e., vegetative cells in log-phase and resting cysts) and grown at two temperatures (18°C and 28°C), plus *Euplotes vannus* and *Strombidium sulcatum* at log phase grown at four temperature treatments (i.e., 16°C, 21°C, 25°C and 16*°C; 9). The genomic DNA of cDNA samples was completely digested at 37ºC using a TURBO DNA-free Reagent Kit (Invitrogen), in which the digestion mixture consisted of 1 μl cell lysate, 1 μl 10×TURBO DNase buffer, 1 μl TURBO DNase (2 units μl ^-1^), and 7 μl nuclease-free water. The optimal DNase dose (2 units) and incubation time (1 h) for DNA degradation were achieved by trail tests, and complete degradation was verified by PCR amplification of 35 cycles and absence of target bands on 1.5 % agarose electrophoresis gel (Fig. S2).

The V4 hypervariable region of 18S rDNA and cDNA in *C. steinii* and *C. inflata* (373 and 374 bp in length, respectively) was amplified using the eukaryotic specific primers TAReuk454FWD1 (5’-CCAGCASCYGCGGTAATTCC-3’), TAReukREV3 (5’-ACTTTCGTTCTTGATYRA-3’) (37). The V1-V3 regions of 18S rDNA and cDNA in *E. vannus* and *S. sulcatum* (445 and 446 bp in length) were amplified with primers Euk82F (5’-GAADCTGYGAAYGGCTC-3’) and Euk516R (5’-ACCAGACTTGCCCTCC-3’) (38, 39). A 6-bp sample identifying barcode was added to both primer sequences. The PCR reaction solution (30 μL) contained 10 ng of DNA or cDNA, 0.2 μM of each primer, and 15 μL of 2× Phusion High-Fidelity PCR Master Mix (New England Biolabs). All PCR amplification reactions were carried out on a T100 Thermal Cycle (Bio-Rad) as follows: initial pre-denaturation at 98°C for 1 min, followed by 30 cycles of denaturation at 98°C for 10 s, annealing at 50°C (56°C for the two marine ciliates) for 30 s, and elongation at 72°C for 30 s, with a final extension step at 72°C for 5 min. The libraries were prepared using a NEB Next Ultra DNA Library Prep kit. Paired-end 250 bp and 300 bp sequencing was executed on HiSeq and MiSeq platforms (Illumnina, USA) for the soil and marine species, respectively. A total of 80 single-cell rDNA and rRNA pools of the two *Colpoda* species were sequenced. For each *Colpoda*, 20 cells were used to sequence DNA and another 20 for cDNA. There were five biological replicates for each treatment. Four samples failed to be amplified, resulting in 39 samples of cells at log phase and 37 samples of resting cysts. A total of 48 single cells of *E. vannus* and *S. sulcatum* were also sequenced. For each species, 24 cells were used to sequence DNA and 24 for RNA pools. Each treatment was run in triplicate.

To assess possible experimental errors during PCR amplification and high throughput sequencing, four clone libraries of the 18S rDNA and cDNA (rRNA) amplicons for the four species were constructed as previously described (19). Five transformed clones were randomly and individually selected from each library and sequenced using MiSeq, as described above. Our expectations were that: (1) the resulting reads of each species would be identical; any observed variants would represent sequencing errors; and (2) only a single unique sequence would be retained for each species after sequencing errors were filtered out.

### Sequence data processing and analysis

The rDNA and cDNA pools of the four ciliate species yielded a total of 8,478,407 reads (Table S1). The primers were removed from the raw reads using Cutadapt v1.18 (29). The DADA2 and DECIPHER package v1.14.1 (40) was applied to model and correct substitution errors, and filter and cluster the amplicons in R (v3.6.3; 41). Reads were trimmed and filtered using the command “filterAndTrim” with the following parameters: minLen=180, maxEE = c(2,2), maxN = 0, truncQ = 2, rm.phix = TRUE, multithread = TRUE. The filtered sequences were dereplicated using the function “derepFastq” to generate unique sequences. The error models were trained using the function “learnErrors” and used for sample inference of the dereplicated reads using the core function “dada” with pool argument set to pseudo, to generate amplicon sequence variants (ASVs). Afterwards, forward and reverse reads were merged using the command “mergePairs”, and “removeBimeraDenovo” was used to check for chimeras. The “IdTaxa” command of package DECIPHER v2.14.0 (27) was used for taxonomic assignment using the SILVA_SSU_v138 training set as the reference database, and the reads assigned to each species were extracted using tool *seqtk* (https://github.com/lh3/seqtk). The “deunique.seqs” command in Mothur was used to create a redundant fasta file from the ASVs fasta and sequence count file. For the treatments of each species, the redundant ASVs were further clustered into operational taxonomic units (OTUs) at a series of similarity thresholds ranging from 89% to 100% using UCLUST.

### Statistical analysis

One-way ANOVA with LSD post hoc test was performed to examine the significance (*α =* 0.05) of the growth phase-wise differences in phenotypic (i.e., cell volume, macronuclear volume, cell to macronuclear volume ratio) and ribotypic traits (per-cell rDNA and rRNA CNs and concentrations, rDNA: rRNA CN ratio). The maximum growth rates of two *Colpoda* species at two different temperatures were statistically compared using *t*-test. Pearson correlation and linear regression analyses were carried out to explore associations between phenotypic and ribotypic traits or between ribotypic traits. All analyses were executed using the statistical software SPSS v.13.0 (SPSS, Chicago, IL).

### Data Availability

The reads from the high throughput sequencing of eukaryotic 18S rRNA genes and transcripts are available under accession numbers SRR10589967 - SRR10590042 (*Colpoda steinii* and *Colpoda inflata*) and SRR10590394 - SRR10590441 (*Euplotes vannus* and *Strombidium sulcatum*). The reads obtained from individual clone sequencing are available under the BioProject accession number PRJNA737167, with accession numbers SRR14800224 - SRR14800243.

## Results

### Cell volume, macronuclear volume, and nucleocytoplasmic ratio

The lag, log, plateau and resting cyst phases of *Colpoda steinii* and *C. inflata* were distinguishable when cultured under replete bacterial prey conditions at both 18°C and 28°C (Fig. 1A). At 28°C, the two species grew approximately 2 and 6 times faster than at the 18°C treatments, respectively. At both temperatures, the smaller species (*C. steinii*) grew consistently faster than the larger one (*C. inflata*): 7.2 ± 0.87 d^-1^ (mean ± SE) vs. 5.1 ± 0.09 d ^-1^ at 28°C, and 3.6 ± 0.32 d ^-1^ vs. 0.9 ± 0.32 d ^-1^ at 18°C (Fig. 1B). Abundant unstable cysts were successfully obtained by chilling actively growing cells of *C. steinii* at 0°C for 4 h and *C. inflata* at 4°C for 10 h (Fig. S3).

The cell volume (CV) of both *Colpoda* species was progressively reduced through lag, log, and plateau to resting cyst phases at both temperatures (Figs 1C, D and 2A). In the 18°C treatment of *C. inflata*, the largest CV (about 2.14 × 10^5^ μm ^3^) was recorded at the lag phase. Upon entering the exponential phase, cells became smaller, and further shrank by 46% during the plateau phase. Relative to the log-phase cells, the resting cysts of this species were greatly reduced in size by 83%. *Colpoda inflata* cells were significantly smaller in the 28°C versus the 18°C treatment (*P* < 0.05 in all cases; Fig. 2A). Compared to the log phase cells, the CV of the small-sized *C. steinii* was reduced by 85% and 69% during encystment at 18°C and 28°C treatments, respectively. At any given phase, the CV was always larger in the cultures reared at 18°C rather than at 28°C (*P* < 0.05), except for the resting cysts, whose size (about 2.0 × 10^3^ μm^3^) was not significantly different between these two temperature treatments (Fig. 2A).

**Figure 2.**
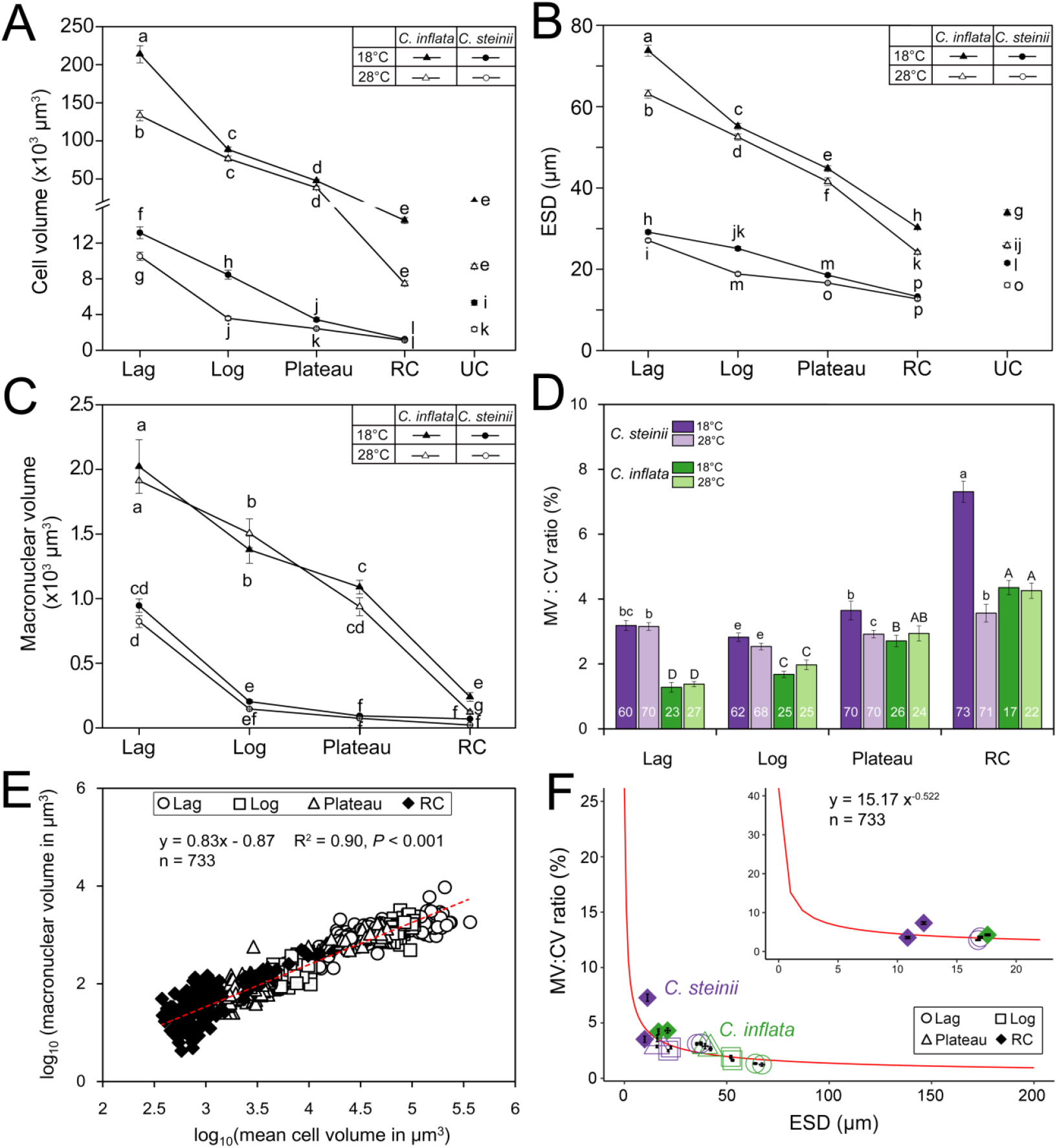
Variation in phenotypic traits of *Colpoda steinii* and *C. inflata* across life-cycle stages and at two temperatures. Unconnected points in line graph panels indicate unstable cysts. (**A**) Cell volume progressively decreased from lag to resting cyst phases. In general, unstable cysts were larger than resting cysts. Cells at a given growth phase were consistently larger at 18°C than at 28°C. (**B**) Equivalent spherical diameter (ESD) progressively decreased from lag to resting cyst phases. **(C)** Macronuclear volume changed similarly to cell volume across stages, but was mostly not significant between the two temperatures. **(D)** Ratio of macronuclear to cell volume (MV:CV) across life-cycle stages. Numbers in the bars indicate the numbers of celled counted. **(E)** A linear log-log relationship between macronuclear (MV) and cell volumes (CV) across life-cycle stages (lag, log, plateau and RC). **(F)** A modeled scaling relationship between MV:CV ratio and equivalent spherical diameter (ESD). Nucleocytoplasmic ratio varies dramatically in small cells [e.g., pico- (< 2 μm) and nano-sized protists (2 ∼ 20 μm)], but changes only slightly in large cells [e.g., microplanktons (20 ∼ 200 μm)]. Non-sharing of common letters indicates significant differences (*P* < 0.05). RC = resting cyst; UC = unstable cyst.

The unstable cysts of both *Colpoda* species were generally larger than the resting cysts, but smaller than the plateau phase cells (Fig. 2A). At 18°C, the volumes of unstable cysts of *C. steinii* and *C. inflata* were on average 5.3 × 10^3^ and 0.21 × 10^5^ μm^3^, respectively (Figs 1C and 2A). A similar pattern of unstable cyst size was observed at 28°C (Fig. 2A). Assuming a globular body shape for *C. steinii* and *C. inflata* cells, the equivalent spherical diameter (ESD) of the cells ranged from 13 to 29 μm and from 25 to 74 μm, respectively, with cross -phase patterns similar to those of CV (Fig. 2B).

Macronuclear volume (MV) exhibited a similar trend to the CV in both *Colpoda* species, i.e., decreasing gradually through lag, log, and plateau phases to resting cyst, regardless of cultivation temperatures (Fig. 2C). *Colpoda inflata* had a consistently larger MV than *C. steinii* at any given phase (*P* < 0.05). For both species, when the MV was relatively smaller (e.g., MV < 500 μm ^3^, or macronuclear diameter < 10 μm), it was consistently and significantly reduced in the 18°C treatments relative to the 28°C treatments. However, when the macronucleus was larger (i.e., macronuclear diameter > 10 μm), the effect of temperature increase on MV was insignificant (*P* > 0.05; Fig. 2C).

The nucleocytoplasmic ratio of two *Colpoda* species ranged from approximately 1% to 8%, and tended to progressively increase from lag, log, to plateau phases and resting cysts (Fig. 2D). The log-phase cells of *C. steinii* had the lowest nucleocytoplasmic ratios (on average 2.92% to 2.61% at 18°C and 28°C, respectively), which were 1.13 to 1.32 times lower than these in lag and plateau phases, respectively. Culturing at the higher temperature generally did not induce significant decreases in the nucleocytoplasmic ratio in most growth phases of either species (*P* > 0.05), except for resting cysts of *C. steinii*, which had a much higher nucleocytoplasmic ratio (on average 7.93%) at 18°C than that at 28°C (3.82%). Compared with the small species, *Colpoda inflata* usually had lower cytoplasmic volume ratios in all phases and at both temperatures (Fig. 2D). Regression analysis showed that macronuclear volume scaled well with *CV*^0.83^, and the nucleocytoplasmic ratio fit well to a power-lower function of *ESD*^-0.522^ (Fig. 2E, F).

### Variations in single-cell rDNA and rRNA copy numbers

The per-cell rDNA and rRNA copy numbers varied significantly among growth phases (ANOVA, *P* < 0.001), showing a consistently decreasing trend from the lag, log, and plateau phases to the resting cyst stage, for both *Colpoda* species and at both temperatures. The resting cysts of both *Colpoda* species lost 50% ∼ 90% of rDNA and over 89% ∼ 99.99% of rRNA copies relatively to the cells in log phase (Fig. 3A, B). The rDNA amount was reduced greatly in the resting cysts of this species at 18°C to about 1000 copies per cyst (Fig. 3A). Although the unstable cysts of *C. steinii* were smaller than the plateau-phase cells, the two had a similar copy number of rDNA [(2.0 ± 0.2) × 10 ^3^]. Compared with the lower temperature treatments, *C. steinii* cultured at 28°C had lower per-cell rDNA copy numbers, particularly for the cells at log and plateau phases (*P* < 0.05).

**Figure 3.**
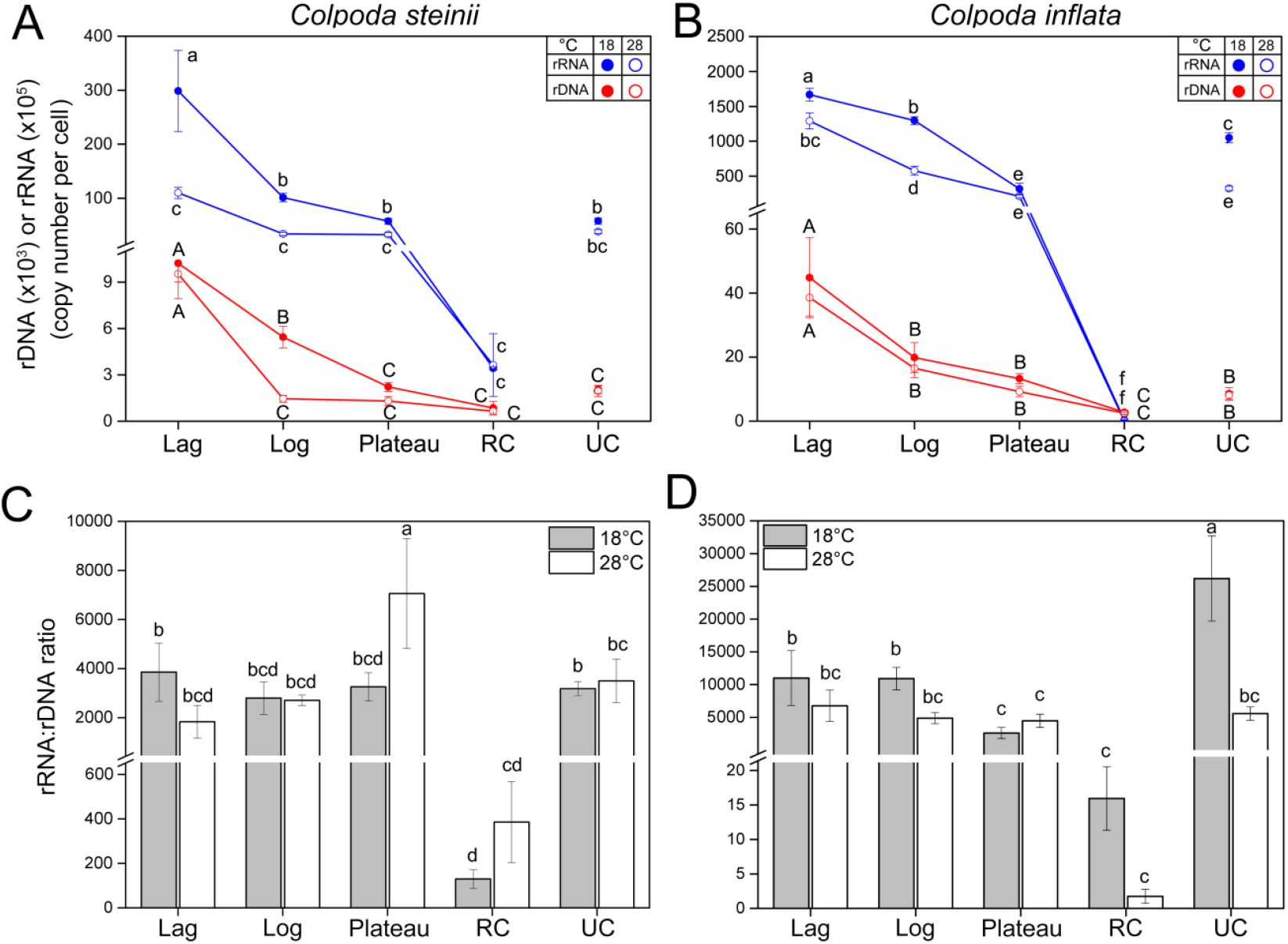
Variation in ribotypic traits of *Colpoda steinii* (A and C) and *C. inflata* (B and D) across lag, log, plateau, resting (RC) and unstable cyst (UC) stages and at two culturing temperatures. Unconnected points in line graph panels indicate unstable cysts. (**A, B**) Single-cell 18S rDNA and rRNA copy numbers progressively and significantly decreased from lag to RC phase. Cells in the UC stage generally had similar amounts of rDNA and rRNA as the vegetative cells in plateau phase. (**C, D**) Copy number ratios of rRNA to rDNA in single cells were significantly lower in resting cysts of both species. The ratio was consistent in all other life-cycle phases of *C. steinii*, and lower in plateau and unstable cystic phases of *C. inflata* (relative to its log phase). Bars represent standard errors. Treatments not sharing common letters indicate significant differences (*P* < 0.05).

The large-sized *Colpoda* species had many more rDNA copies than the small-sized congener across all life-cycle stages (e.g., 1.98 × 10^4^ vs. 5.4× 10^3^ at log phase; Fig. 3B). Phase-specific comparisons showed that warming had no significant effect in modulating per-cell rDNA content in this species (*P* > 0.05). The resting and unstable cysts of *C. inflata* contained about 2500 and 8100 rDNA copies, respectively, which were about 3.3 and 4.2 times higher than those of *C. steinii* (Fig. 3B).

The cellular rRNA copies were far more abundant, and varied more dramatically across growth stages, than the rDNA copies in both *Colpoda* spp. (Fig. 3A, B). In the 18°C treatment, a single cell of *C. steinii* had 3.0 × 10^7^, 1.0 × 10^7^, 5.7 × 10^6^, and 3.4 × 10^4^ rRNA copies at the lag, log, plateau and resting cyst stages, respectively (Fig. 3A). At the same temperature, the rRNA molecules were much more abundant in *C. inflata*, with 1.7 × 10^8^, 1.3 × 10^8^, 3.2 × 10^7^ and 3.1 × 10^4^ copies across these phases (Fig. 3B). Although smaller than the plateau-phase cells, the unstable cysts had relatively similar (in *C. steinii*) or even higher number of rRNA copies (in *C. inflata*) (Fig. 3A, B). Comparing the two temperature treatments, warming always led to a significant decrease in cellular rRNA amount of both species and across all stages plus unstable cysts (*P* < 0.05). A cell at any vegetative growth phase of these two species had approximately 200 to 5000 times more rRNA copies than resting cysts (Fig. 3A, B).

Differences between the two *Colpoda* species in terms of the patterns of rRNA:rDNA CN ratio across growth phases and between temperature treatments were minor (Fig. 3C, D). The rRNA: rDNA CN ratios in *C. steinii* ranged from 2000 to 4000, without statistical differences among the lag, log, plateau and unstable cystic stages (*P* > 0.05; Fig. 3C). Nevertheless, at 18°C the rRNA: rDNA ratios in *C. inflata* decreased from 10000 to 2000 when population growth entered the plateau phase; the rRNA: rDNA ratio of unstable cysts was also significantly high in the 18°C treatment in this species (Fig. 3D). Furthermore, while the rRNA: rDNA ratios in log-phase cells, unstable, and resting cysts of *C. inflata* significantly dropped at the higher culturing temperature (Fig. 3D), the same was not observed in *C. steinii* (Fig. 3C). Nevertheless, the resting cysts of both species had much lower rRNA: rDNA ratios (150 – 400 in *C. steinii*, and 2 – 16 in *C. inflata*) than the cells in all other phases (Fig. 3C, D).

### Linking ribotypic with phenotypic traits

Based on all CN data of vegetative and cystic cells of these two *Colpoda* spp., correlation and linear regression analyses showed that both rDNA (*rDNA*_*CNPC*_) and rRNA copy number per cell (*rRNA*_*CNPC*_) were positively and significantly related to CV or ESD, and could fit well to linear regression curves (*R*^2^ = 0.91, *P* <0.01; Fig. 4A; Table 1, Eq. 1 and 2), irrespective of cell cycle stage.

**Figure 4.**
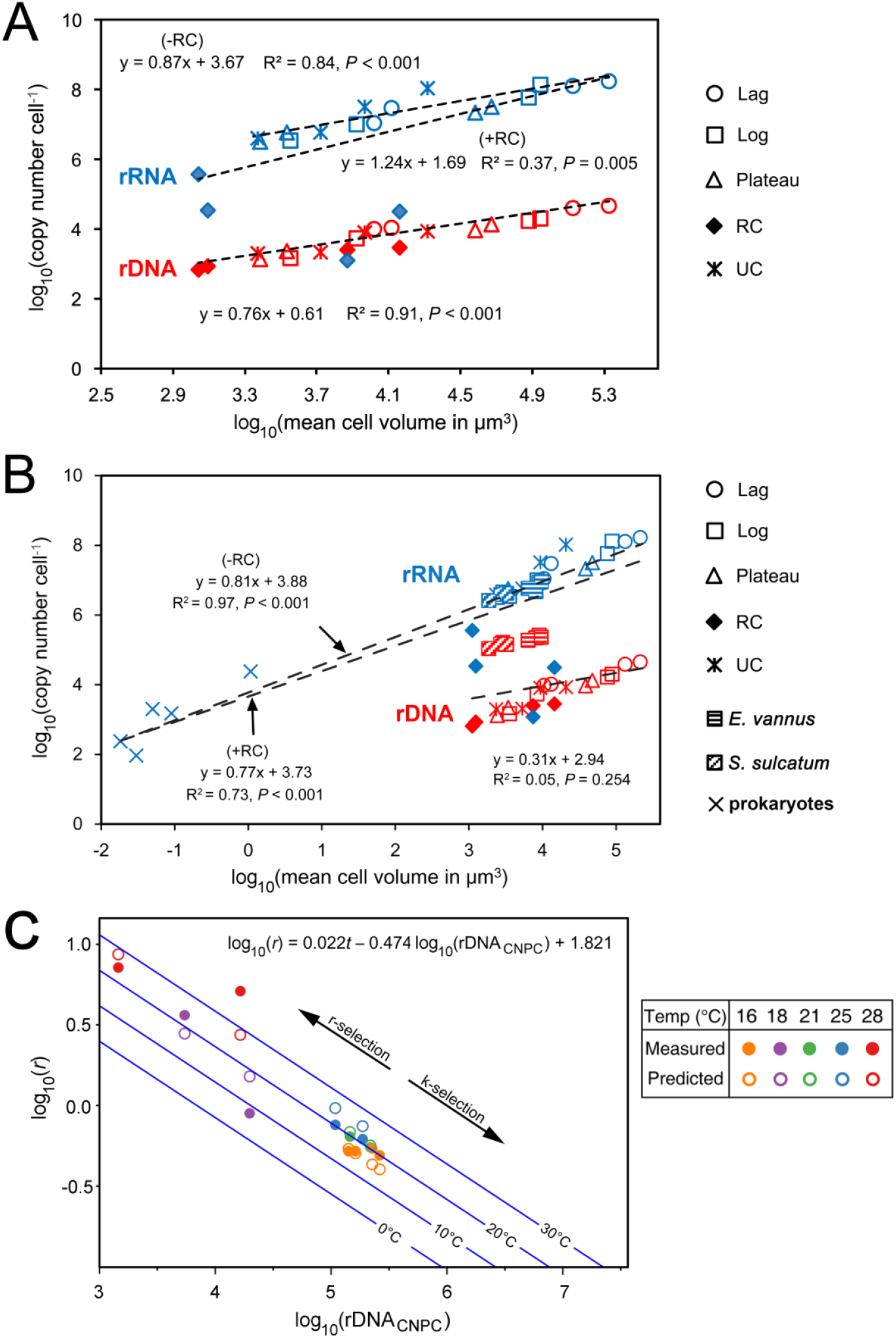
Scaling relationships among ribotypic traits, growth rate, and cellular attributes. **(A)** The per-cell rDNA and rRNA copy numbers (CNs) in two *Colpoda* species across life-cycle stages was significantly related to cell volume. **(B)** Scaling relationships based on a combined dataset consisting of two *Colpoda* species (present study), *Euplotes vannus* and *Strombidium sulcatum*, and five prokaryotic species (Fu and Gong 2017). The per-cell rRNA CNs are better predicted using cell volume when resting cysts (indicated by four symbols of solid diamond in blue) are excluded from the regression analysis (-RC) than the resting cystic data are incorporated (+RC). The rDNA CNs of *E. vannus* and *S. sulcatum* were much higher than those of *Colpoda* cells with similar size ranges, suggesting that rDNA CN and CV relationship may be conserved among closely related taxa, but becomes inconsistent among distant taxa. (**C**) Maximum growth rates (*r*) of *Colpoda steinii, C. inflata, E. vannus* and *S. sulcatum* can be well predicted by rDNA CN per cell (CNPC) and temperature (*t*). Lines in blue denote predicted growth rate - CNPC relationships at 0, 10, 20 and 30°C. The amount of rDNA copies in genomes is inversely related to the growth potential of the ce ll, which might underline r- and K-selection of these organisms. RC = resting cysts; UC = unstable cysts.

**Table 1.**
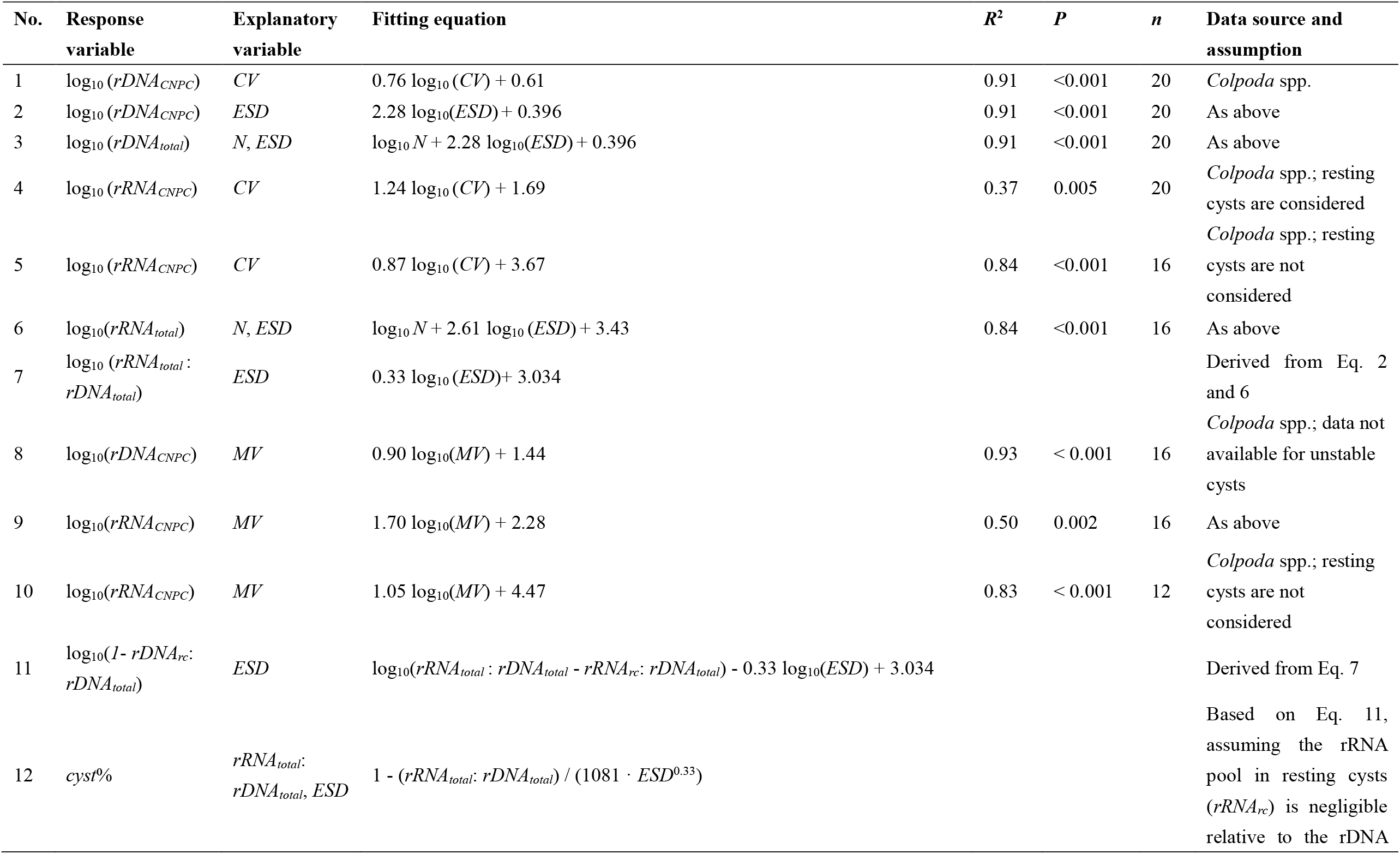
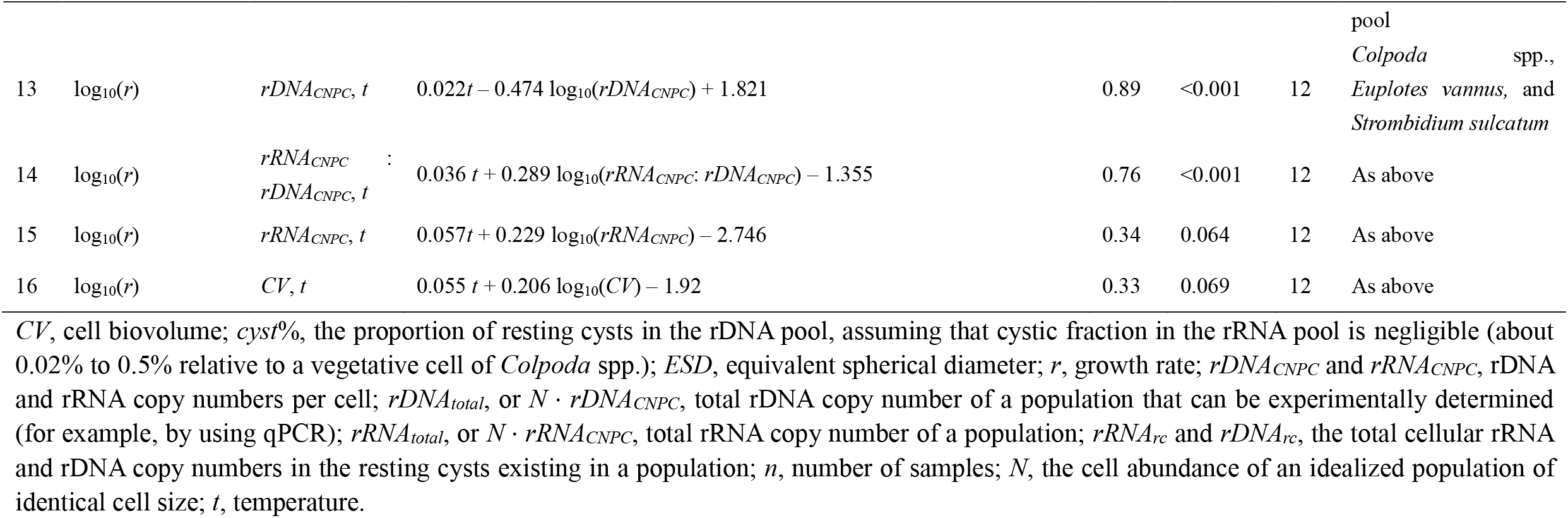
Fitting equations linking ribotypic traits with phenotypic traits in ciliated protists

For an idealized population of similar cell size, the total rDNA copy number of the population (*rDNA*_*total*_ or *N* · *rDNA*_*CNPC*_) can be expressed as a function of *ESD* and abundance of the cells (*N*) (Table 1, equation 3). Since *N* · *rDNA*_*CNPC*_ can be experimentally determined (for example, by using qPCR), this equation indicates that inferring cell abundance of a population using rDNA copy numbers depends on cell size of the population.

The correlation between *rRNA*_*CNPC*_ and CV across all life cycle phases including resting cysts (+RC) was significant (*P* = 0.005; Table 1, equation 4). However, the *R*^2^ score (37%) was only moderate (Fig. 4A). Nevertheless, when the rRNA CN data of the resting cysts (which contained extremely low rRNA amount that could not be easily predicted when using cell size, i.e., the four blue full diamonds that are discrete from other points in Fig. 4A) were excluded from the analysis (-RC), the relationship became well supported (*R*^2^ = 0.84; Table 1, equation 5). Similarly, the total rRNA copy number of the population (*rRNA*_*total*_ or *N* · *rDNA*_*CNPC*_) can be expressed as a function of *ESD* and cell abundance (*N*) (Table 1, equation 6). Based on a combination of equations (3) and (6), which presume all cells in a population are non-resting cysts, the copy number ratio of total rRNA: rDNA depends on a single factor, ESD or cell size (Table 1, equation 7). The correlations between *MV* of cells at all life cycle stages and ribotype CNs exhibited patterns similar to the ones observed for *CV* (see Table 1, equations 8 - 10; Fig. S4).

Assuming that the proportion of resting cysts in the rDNA pool is *cyst*%, and their fraction in the rRNA pool is negligible (about 0.02% to 0.5% relative to a vegetative cell of *Colpoda* spp.), equation (7) can be alternatively expressed as equation (11), then *cyst*% can be assessed using a function of both *rRNA*_*total*_: *rDNA*_*total*_ and cell size (Table 1, equation 12).

According to (9), the per-cell rRNA CNs in the log phase cells of *E. vannus* and *S. sulcatum* were at similar levels as the ones in *Colpoda* cells of similar cell sizes [the lg(*CV*) ranging from 3.3 to 4.1; Fig. 4B]. However, these two marine species had about 30 ∼ 80 times more rDNA copies (109,000 ∼ 263,000 per cell) than *C. steinii* cells (1300 ∼ 9540 copies per cell) within the same cell size range (Fig. 4B). In other words, the previously data on *E. vannus* and *S. sulcatum* largely followed the scaling relationship between rRNA and CV of non-resting *Colpoda* (*R*^*2*^ = 0.86, *P* < 0.001) described herein, but was not consistent with the rDNA-CV relationship described in the present study (*R*^*2*^ = 0.05, *P* < 0.001).

Interestingly, based on a combination of datasets comprising the previously reported for *Euplotes vannus* and *Strombidium sulcatum* (*n* = 8; 9) along with the present data for *Colpoda* spp. (*n* = 4), the maximum growth rate (*r*) of these species at a given culturing temperature (*t*) can be well predicted by *rDNA*_*CNPC*_ (*R*^*2*^ = 0.89, *P* < 0.001, *n* = 11; equation 13; Fig. 4C), and ratio of *rRNA*_*CNPC*_ : *rDNA*_*CNPC*_ (*R*^*2*^ = 0.76, *P* < 0.001, *n* = 12; equation 14), but poorly by *rRNA*_*CNPC*_ (*R*^*2*^ = 0.34, *P* = 0.064, *n* = 12; equation 15), or CV (*R*^*2*^ = 0.33, *P* = 0.069, *n* = 12; equation 16).

### Single-cell landscape of rDNA and rRNA variants

Processing the high throughput sequencing data of 18S rDNA and cDNA (rRNA) amplicons of individual clones indicates that application of the DADA2 pipeline effectively filtered out error ASVs. Consequently, a single ASV was obtained for *E. vannus* and *S. sulcatum*, and 12 ASVs for each of the two *Colpoda* species (Table S2). For the latter two species, ASV clones with sequence divergence lower than 1% were lumped into a single OTU at a similarity cutoff of 99% (OTU_99%_) to offset the sequencing errors, in order to examine the biological variations by counting OTU numbers (Table S2). In the following sections, for *E. vannus* and *S. sulcatum*, once multiple ASVs (equivalent to OTUs clustered at a 100% similarity level, i.e., OTU_100%_) and multiple OTU_99%_ were detected, they were considered to reflect real biological polymorphisms of rDNA and rRNA.

A total of 123 rDNA and cDNA samples derived from 123 single cells of the four ciliate species were successfully sequenced. For all DADA2-filtered sequences of a single cell, one to multiple OTU_99%_ were often detected in rDNA and rRNA pools: in *C. steinii*, 1 - 5 OTUs; and in *C. inflata*, 1 - 4 OTUs. For OTU_100%_, in *E. vannus* 1 - 6 OTUs were detected and in *S. sulcatum* 2 - 6 OTUs (Table S1). In each species, there was a single, numerically dominant OTU, the relative abundance of which varied widely (87% - 100%), depending on species, rDNA or rRNA pools, life cycle stage, and temperature (Fig. 5A-D). Significant changes in the proportion of dominant OTUs were detected in rRNA pools, but not in rDNA (Fig. 5A-D). Furthermore, the dominant rRNA-based OTU_99%_ was present in significantly lower proportions in the resting cysts of *C. steinii* (at 28°C) and *C. inflata* (at 18°C) than in log phase *Colpoda* cells at the same temperatures (Fig. 5A, B). Temperature generally did not affect the proportion of dominant OTU in any species (ANOVA, *P* > 0.05; Fig. 5A-D), except for two pairwise comparisons in the case of *E. vannus*, in which the dominant OTU_100%_ of rRNA was present in a significantly lower proportion in the touchdown treatment (98.47 ± 0.06%) than in the treatments at 16°C and 21°C (Fig. 5C).

**Figure 5.**
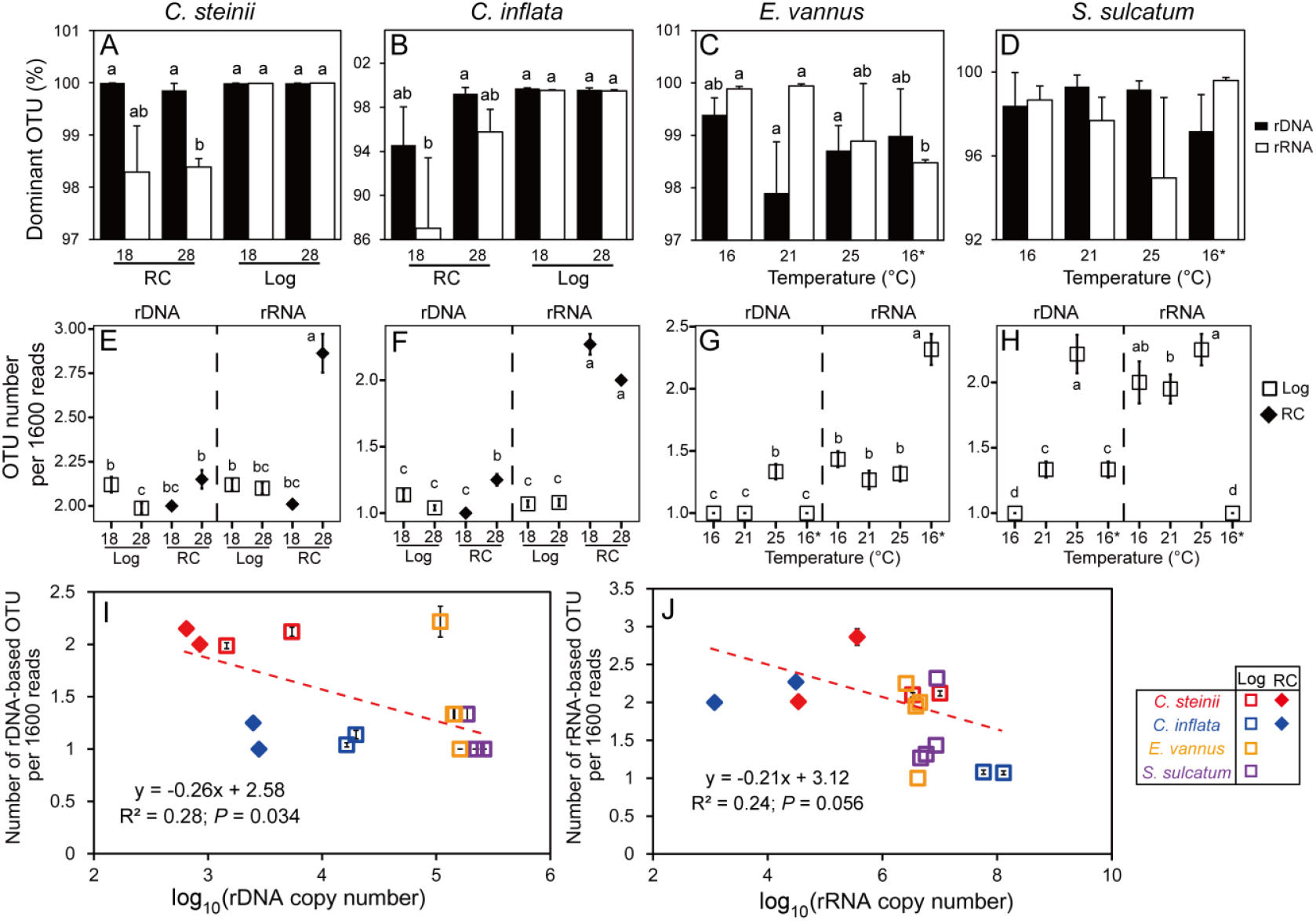
Characterization of operational taxonomic units (OTUs) in rDNA and rRNA pools in single cells of four ciliate species. **(A - H)** Comparisons of the relative abundance of the most dominant OTU (**A - D**) and OTU numbers per 1600 reads (**E - H**) between resting cysts (RC) and log-phase vegetative cells (Log) of *Colpoda steinii* and *C. inflata* at 18°C and 28°C, and in a single log-phase cell of *Euplotes vannus* and *Strombidium sulcatum* at four temperatures. **(I, J)** Both rDNA (**I**) and rRNA-based OTU numbers (**J**) are negatively related to sequence number within a single cell. Two treatments not sharing common letters indicate significant differences (multiple pairwise comparisons, *P* < 0.05). Note that the OTUs are defined at a sequence similarity cutoff of 99% for *C. steinii* and *C. inflata*, and at 100% for *E. vannus* and *S. sulcatum*, based on the results of sequencing and analysis of individual clones with 18S rDNA or cDNA fragments.

The numbers of rDNA- and rRNA-based OTUs per 1600 reads in single cells varied according to life cycle stage and temperature (Fig. 5E-H). At 28°C, resting cysts consistently had higher numbers of OTU_99%_ than log phase cells in both *Colpoda* species (*P* < 0.05; Fig. 5E, F). The log vs. resting phase cells of *C. steinii* at higher temperature had fewer rDNA-OTU_99%_, while the resting vs. log phase cells of *C. inflata* had more (*P* < 0.05; Fig. 5E, F). For *E. vannus* and *S. sulcatum*, the highest numbers of rDNA-based OTU_100%_ were consistently detected at 25°C (*P* < 0.05, Fig. 5G, H). Nevertheless, the number of rRNA-based OTU_100%_ peaked at 25°C in *S. sulcatum* and at 16°C in *E. vannus* (*P* < 0.05; Fig. 5G, H). Regression analysis based on single-cell data of all four species indicated that the number of OTUs in the rDNA pool was significantly and negatively related to cellular rDNA CN (*R*^*2*^ = 0.28, *P* = 0.034; Fig. 5I), while the number of OTUs in rRNA pool was marginally and negatively related to cellular rRNA CN (*R*^*2*^ = 0.24, *P* = 0.056; Fig. 5J). However, analogous relationships between rDNA and rRNA OTU numbers and cell volume were not significant (*P* > 0.66; Fig. S5).

### Divergence in rDNA and rRNA sequences in a single cell and its effect on OTU clustering

By clustering all the rDNA variants obtained from single-cell samples into OTUs at a series of gradually lower cutoffs of sequence similarity, the numbers of identified OTUs generally decreased (Fig. 6A). At the 97% cutoff, an average of 2.0 - 3.0 rDNA-based OTUs remained for *C. steinii*, and 1.0 - 2.0 for *C. inflata*. The observed OTU numbers leveled off to 1 (or insignificantly different from 1) when the cutoff decreased to 89% in *C. steinii* and 95% in *C. inflata* (Fig. 6A). A single DNA-based OTU was identified for *E. vannus* and *S. sulcatum* at the 99% cutoff. Taking the sequencing errors that remained after DADA2 filtering into account (1% in the two *Colpoda* species and nil in *E. vannus* and *S. sulcatum*; Table S2), it is estimated that the real maximum differences among the rDNA variants in single cells of these species were 10%, 4%, 1% and 1%, for *C. steini, C. inflata, E. vannus* and *S. sulcatum*, respectively.

**Figure 6.**
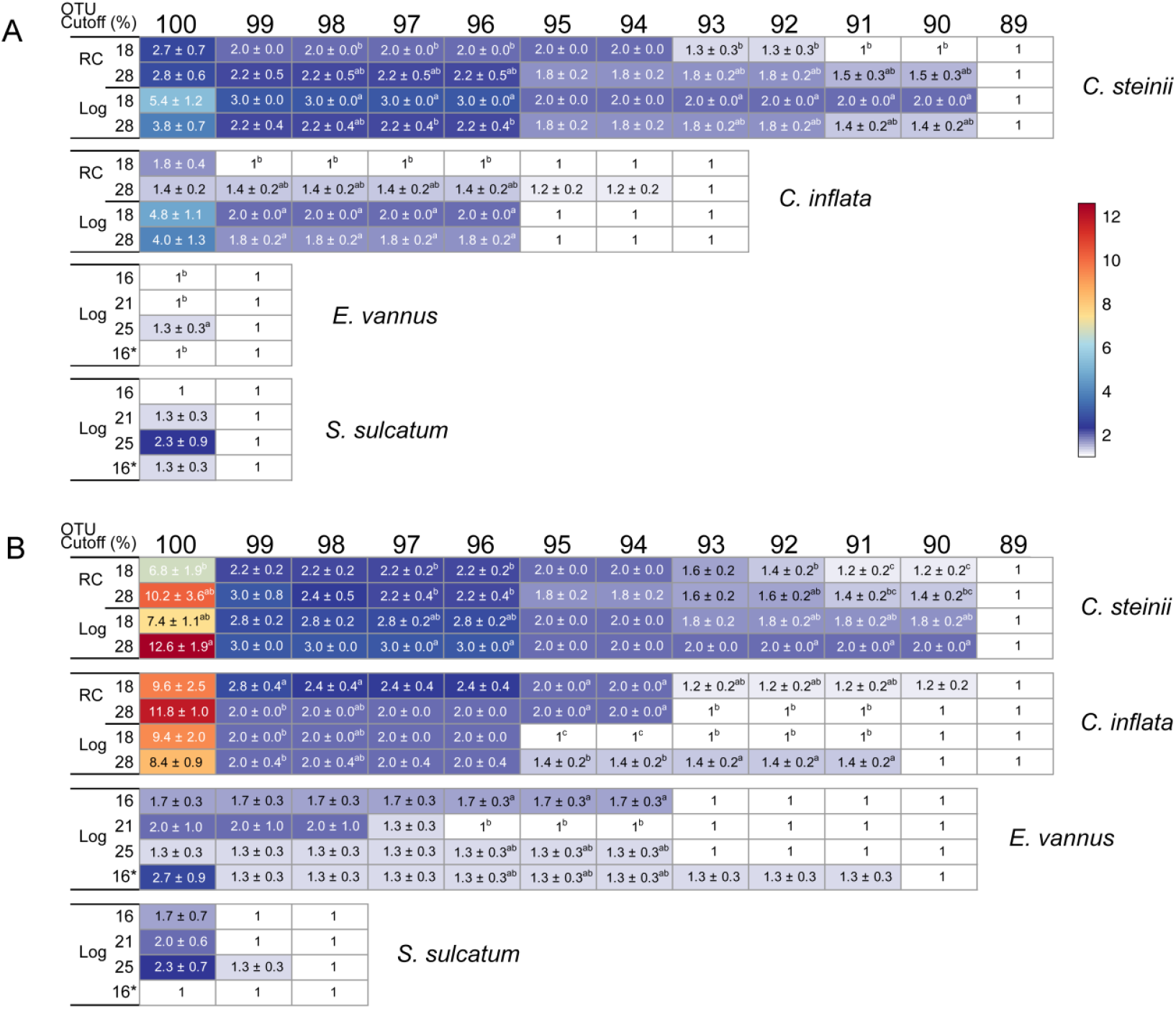
Variation in the numbers of OTUs in rDNA (A) and rRNA (B) pools of single cells of ciliated protists. The OTUs were defined at a sequence similarity threshold ranging from 89% to 100%. The nominal maximum intra-individual rDNA and rRNA sequence differences of the four species ranged from 1% (in *S. sulcatum*) to 11% (in *C. steinii*). Two treatments not sharing common letters indicate significant differences (multiple pairwise comparisons, *P* < 0.05). The color bar indicate number of OTUs per cell.

For all species, the sequence divergence of rRNA molecules was generally higher than that of rDNA (Fig. 6B). *Colpoda steinii, C. inflata* and *E. vannus* had multiple (2.2 - 3.0, 2.0 - 2.4, and 1.3 - 1.7) OTUs at the 97% cutoff, but only 1 OTU when the cutoff was raised to 89%, 90%, and 93%, respectively. Nevertheless, the number of rRNA OTUs in *S. sulcatum* was rapidly reduced at the 99% cutoff. The differences in rRNA sequences are estimated up to 10%, 9%, 7% and 1% in *Colpoda steinii, C. inflata, E. vannus*, and *S. sulcatum*, respectively (Fig. 6B). At the 97% cutoff, significantly different rDNA-based OTU numbers were frequently identified between resting and log phase cells of *Colpoda* spp. (Fig. 6A), however this was never the case between any two temperature treatments in any species (Fig. 6A, B).

## Discussion

### Ribotype copy number variations across different phases of growth and cystic forms

Substantial variations in rDNA and/or rRNA quantities have been previously recorded in a range of protistan species at various phases of growth or physiological states (e.g., a *Tetrahymena* ciliate, 42; a raphidophyte, 43; a foraminifer strain, 10; and dinoflagellates, 44-46). Nevertheless, this study provides the first evidence of linkage between ribotypic and phenotypic traits in ciliated protists. Specifically, the per-cell CN and cellular concentrations of rRNA and rDNA molecules, along with their CN ratio were quantitatively linked with cell and macronuclear sizes. This was a consistently observed relationship across all growing phases and dormant cystic forms.

Artificially induced unstable cysts constituted an exception to this observation. The per-cell rDNA and rRNA CNs, as well as, cell size (Figs 2A, B, 3A, B) were significantly reduced in these cysts. Freezing raw environmental samples (e.g. soil, sediment or water) temporarily to preserve microbiomes is occasionally used in molecular ecology. Herein, we show that this practice might artificially alter both phenotypic and ribotypic traits of cyst-forming protists, leading to underestimation of their biomass and proportions of rDNA and rRNA quantities in the ribotypic pools. Nevertheless, chilling treatment is a simple, non-destructive approach that arrests growing cells at the unstable cyst stage, thus narrowing down the cell size spectrum of a population consisting of different-sized cells in various growth phases, while preserving cell numbers in the original samples. Thus, chilling treatment has the potential to enhance the accuracy in estimating population abundances.

We provide evidence that both unstable and resting cysts of *Colpoda* follow the rDNA CN-CV relationship (Fig. 4A; Equations 1 and 2), so their biomass can be reasonably predicted in rDNA-based molecular surveys. Nevertheless, compared to unstable cysts, resting cysts have disproportionately fewer rRNA molecules, which could be attributed to the former having a higher metabolic rate than the latter (47). Based on the ribotypic distinctness of resting cysts, we propose a theoretical framework to estimate their proportion in the rDNA pool of a population using experimentally determined rRNA:rDNA ratio of a given *Colpoda* species. The proposed framework supports the use of rRNA:rDNA ratio as a predictor of protistan dormancy in environmental studies. Importantly, the framework emphasizes taking cell size into account (see Equation 12), which is especially relevant when comparing cellular activity across species or taxa of contrasting cell sizes. Thus, species- or taxon-specific qPCR assays could be applied in future studies for quantifying cysts of *Colpoda*, and/or other protists in environmental samples.

### Cellular rRNA content scales inversely with cell size, but maximum growth rate depends on rDNA content

Previously, a scaling relationship between rRNA content and cell size of log-phase cells of two protist species and five species of bacteria and archaea was proposed (9). Herein, we show that the scaling relationship holds across life cycle stages, indicating that the scaling law between rRNA content and cell/body size applies not only to macro-organisms, but microbes as well (48). Nevertheless, the scaling relationship did not apply to resting cysts, as their rRNA content was significantly lower than expected given their cell size. This indicates that rRNA quantity is a suitable molecular indicator for providing a biomass-based view of microbial community structure. In support of this, previous studies of mock protistan communities showed consistent patterns when using rRNA and cell counting approaches (49).

Unlike rRNA, the scaling relationship between rDNA and cell volume was not uniform for any of the four ciliate species examined. Instead, two separate functions were applied; one for *Euplotes* and *Strombidium* (rDNA ∼ CV^0.44^; 9) and another for *Colpoda* spp. (rDNA ∼ CV^0.76^; this study). This is primarily due to the much higher cellular rDNA content in *E. vannus* and *S. sulcatum* (23 - 59 copies per μm^3^) than in the similarly sized cells of *Colpoda steinii* and *C. inflata* (0.2 - 0.8 copies per μm ^3^) (Fig. 4B). The disproportion between per-cell rDNA copy number and cell size has been previously pointed out (50). The variable rDNA content of Spirotrichea and Colpodea may stem from their genomic distinctness. Spirotrichean ciliates are well known for their highly polyploidized genomic macronuclear DNA (26). Higher rDNA content also implies higher genomic DNA content and larger macronuclear volumes (51). In support of this, both *E. vannus* and *S. sulcatum* have disproportionately large macronuclei (N/C ratios about 4.71% and 2.73%, respectively) that do not correspond to the predicted values (1.57% and 2.43%) derived from the *Colpoda*-based model (Fig. 2E, F). This indicates that larger macronuclear size does contribute to higher rDNA content in these two spirotricheans. Even when their “enlarged” macronuclear sizes are taken into consideration, their calculated per-cell rDNA copy numbers according to the *Colpoda*-based function (Equation 8) are only about 1/2 and 1/16 of those experimentally determined. This suggests that the high rDNA content of spirotricheans is not just due to their unusual phenotype (enlarged macronuclei), but that certain, as yet undiscovered, mechanisms for genomic innovation may also be at play.

Interestingly, *C. steinii* grew 7 to 12 times faster than *E. vannus* and *S. sulcatum*, despite these three species having comparable cell size and cellular rRNA content. When predicting growth rate (Equations 13 and 15), the model parameterized using rDNA performed much better than the one using rRNA. Thus, besides temperature (which modulates cell size via the temperature-size rule, 52), cellular rDNA content is an important factor in determining generation time of these ciliated protists. This observation suggests that growth rate is not only determined by rRNA content, as proposed in the growth rate hypothesis (53), but can also be modified by rDNA contents. In other words, growth rate depends mainly on rDNA, when the cells have similar cell sizes or rRNA contents (see Fig. 4B).

Existing studies on genomic content and growth rates of various organisms can shed light in understanding the mechanisms underlying modulation of cell growth rate and relationship with rDNA content. For instance, haploid cells of budding yeast grew more rapidly than their diploid counterparts of larger cell size. The observed fitness difference was attributed to their cell surface to volume ratio. Specifically, growth rate was dependent on the transfer of products into cytoplasm generated upon hydrolysis of organic phosphorous by a cell surface-bearing acid phosphatase (54, 55). The surface-to-volume-ratio theory does not seem to apply to phagotrophic ciliates, which graze on food particles through their cytostome and digest them inside cytoplasmic food vacuoles. Nevertheless, the lower nuclear to cell volume ratio of *C. steinii* may allow for a larger cytoplasmic space to accommodate additional food vacuoles, enabling digestion of more bacterial prey and maintenance of a higher rate of material supply for growth than *E. vannus* and *S. sulcatum*. Spirotricheans have a high content of rDNA and genomic DNA, both of which are particularly P- and N-rich. Thus, these ciliates need element-rich resources and invest a longer time for DNA replication, synthesis of associated proteins, and cell division (56). Conversely, the similarly sized colpodean ciliate *C. steinii* with fewer rDNA copies and a more streamlined genome would save more P and N to synthesize rRNA and assemble ribosomes for rapid growth.

### Richness and composition of rDNA and rRNA variants across life cycle stages and at different temperatures

In the present study, the error rate of high throughput sequencing was offset on the basis of individual clones of amplicons. Even so, low to high levels (1% - 10%) of sequence divergence were still identified in single cells, indicating high likelihood of the observed variants of 18S rDNA and rRNA of ciliated protists being of biological origin. The extent of divergence observed in this study is comparable to the previously uncovered using high throughput sequencing (13% in radiolarians, 18; up to about 16% in a *Protocruzia* ciliate, 57). Previous studies on rDNA polymorphisms using clone libraries and Sanger sequencing yielded lower intragenomic divergence (< 2%) in ciliates (9, 21, 50) and diatom species (0.57%-1.81%, 58), while divergence was moderate (about 5%) in foraminifera (17). It has been speculated that use of polymerases of varying fidelity in separate studies might affect polymorphism estimations (21). Herein, identical experiments (e.g., polymerase and PCR thermal cycling conditions) were performed expected to yield similar error rates. This suggests that the observed distinct divergence rates among the four ciliates might be dependent on species identity. A possible reason for this could be that the species with low per-cell rDNA copy numbers tend to have more rDNA variants (Fig. 5I), providing a higher chance to explore cellular rDNA variant space and thus resulting in a higher sequence divergence (Fig. 6A).

The results confirm our expectation that multiple and less abundant (relative abundance ≤ 13%) rRNA variants (or OTU_99%_ of *Colpoda* spp. and OTU_100%_ of *E. vannus* and *S. sulcatum*) are transcriptionally expressed in single cells of the protists studied herein (Fig. 6B). These variants could combine with ribosomal proteins, giving rise to extremely high ribosomal heterogeneity in these ciliates. Such a high diversity of ribosomes in single-celled eukaryotes is surprising, because these organelles have long been thought of as a homogeneous cellular machinery. A notable exception is the parasitic protist, *Plasmodium berghei*, in which two structurally distinct 18S rRNA transcripts were found (22). Recent studies have suggested that ribosomal variants could preferentially bind different mRNAs to regulate gene expression and corresponding phenotypes of bacteria under stress conditions (24, 25, 59). Given the high intracellular rDNA diversity in diverse protistan groups, the significance of ribosomal heterogeneity in the context of physiological and ecological adaptations of protists inhabiting various ecosystems would be an interesting topic to explore.

Pairwise comparisons also support relationships between rDNA- and rRNA-based OTU richness and life cycle stage and temperature (Fig. 5E-H), indicating the dynamic nature of ribotypic composition and its tight coupling with physiological status in ciliates. More specifically, the richness of rRNA-based OTU_99%_ composition in the cysts was higher than that in log phase cells. This could be attributed to shrinkage of the rRNA pool during encystment accompanied by loss of cellular rRNA copies of the dominant variant and maintenance of minor ones (Fig. 5A, B). Functionally, keeping more rRNA OTUs (and by extension, a higher diversity of ribosomes) in the resting cysts may be significant for soil ciliates in maintaining a high metabolic potential to readily respond to favorable environmental conditions. Resting cysts have specialized mRNA and proteins (60), and *C. inflata* is known to undergo demethylation of macronuclear DNA during encystment (61). Further investigations are needed to explore the mechanistic links between DNA methylation, rRNA-derived ribosomal heterogeneity, and mRNA translation towards a better understanding of protistan adaptations and their biological and ecological roles.

### Concluding remarks

In summary, the present study is the first to provide evidence for quantitative relationships between rDNA and rRNA copy numbers and cell size of protists across life cycle stages, illustrate a methodological framework for estimating the quantity of resting cysts, and show compositional changes in sequence variants during formation of resting cysts and in response to temperature. Although the findings on phenotype-ribotype coupling were based on ciliate models in this study, variations in genome content across life-cycle stages and under different conditions have been reported for other protists, e.g., *Giardia* (41) and amoeba species (62). This suggests that relationships among ribotype numbers, genomic content, cell and nuclear size, and growth rate likely exist in many groups of protists, as previously predicted by (32, 51). Our findings have tentatively important implications for microbiome and microbial biogeographic studies of microbial eukaryotes: 1) we demonstrate substantial changes in cellular contents of rDNA and especially rRNA across life-cycle stages of protists. This suggests that, even if the cell abundance of each species in a community remains unchanged, the relative abundances of these molecules among protistan species, and hence the inferred diversity and community structure, can be highly variable depending on life-cycle stage and physiological shifts. This layer of biological complexity should be considered in metabarcoding approaches focused on detecting protistan diversity alterations, ecological responses and adaptations to environmental changes and biotic interactions (e.g., water and prey availability, and warming), thus has functional and ecological significance; 2) cellular rRNA quantity generally scaled much better with cell volume than genomic rDNA, which could be related to genome duplication and reduction, or alteration of ploidy levels among protistan taxa. This implies that rRNA is a better molecular marker than rDNA for mapping biomass partitioning in multi-species communities, especially among active members; 3) in contrast, growth rate of protistan cells could be more accurately predicted using cellular rDNA content rather than rRNA, of rRNA:rDNA copy number ratio, or cell size. Linking this ribotypic trait to cellular growth opens a window for a taxon-specific view of microbial production in various environments; and 4) protistan diversity, community composition, rare-to-abundant transition, seasonality, biogeographical patterns, dynamic responses to perturbations, as well as, ecological and biogeochemical functions in soil, planktonic and benthic systems are all affected by dormancy of cysts (1, 5, 11, 12). Our work illustrates a potential resolution in determining the functional and ecological importance of protistan dormancy across time and space by simultaneously quantifying rDNA and rRNA molecules.

Our results of sequencing and downstream analyses of single-cell rDNA and rRNA variants have implications for ciliate/protistan diversity surveys using metabarcoding: 1) despite using a sensitive pipeline (e.g., DADA2) for quality filtering, low levels of sequencing errors (< 1%) could still exist in 18S rDNA and rRNA amplicon sequences of some species; therefore, it is generally recommended that a 99% sequence similarity threshold be applied if intragenomic polymorphism is of interest. 2) Inferring species-level diversity using a sequence similarity of 97% can lump intragenomic sequence polymorphisms in some, but not all, species. In addressing the changing patterns of protistan richness, this part of richness inflation can be relieved if the inflated species are present in all or most samples or they account for substantially low sequence proportions. Otherwise, the intragenomic polymorphisms of rDNA and rRNA sequences remain an issue that potentially undermines our capability in estimating and predicting protistan richness in natural or complex communities.

## Acknowledgments

This work was supported by the grants from the Marine S & T Fund of Shandong Province for Pilot National Laboratory for Marine Science and Technology (Qingdao) (No. 2018SDKJ0406-4), the Natural Science Foundation of China (Nos. 31572255, 41976128 and 31672251), the Research Fund Program of Guangdong Provincial Key Laboratory of Marine Resources and Coastal Engineering (No. 311021004), and the Key Research Program of Frontier Sciences (QYZDB-SSW-DQC013).

