## Supplemental Files for "Coupling between ribotypic and phenotypic traits of protists across life-cycle stages and temperatures"

**Supporting information**

**Figure S1. Standard curves of quantitative PCR in determining single-cell rDNA or rRNA (cDNA) copy numbers in *Colpoda* spp.** The values of $R^2$ (0.998 ~ 1.000) and amplification efficiencies (E, 98% ~ 108.9%) are shown.

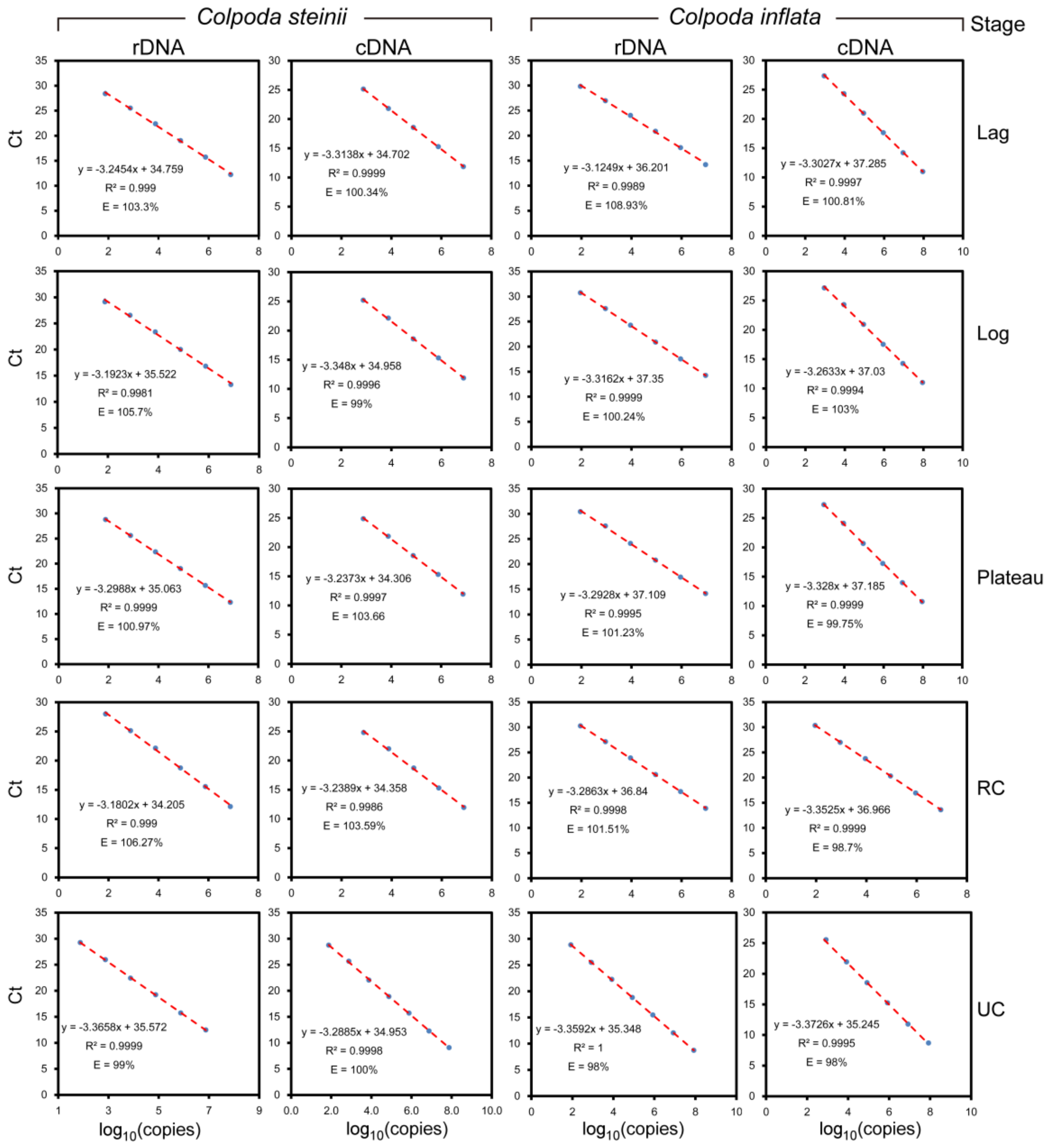

**Figure S2. Electrophoretogram of PCR products obtained via amplification of 35 cycles of rDNA fragment.** The single-cell nucleic acids (containing both rDNA and rRNA) were treated with different amounts of DNase (0 to 3 μL) for various periods of time (10, 30, and 60 min). There were no detectable target bands of rDNA fragments in the samples which were treated with 1 μL (2 U) of DNase for 60 min (lanes 5 and 8), indicating optimal incubation conditions for complete degradation of rDNA. M, DNA markers. Lane 12 indicates negative control with double-distill water.

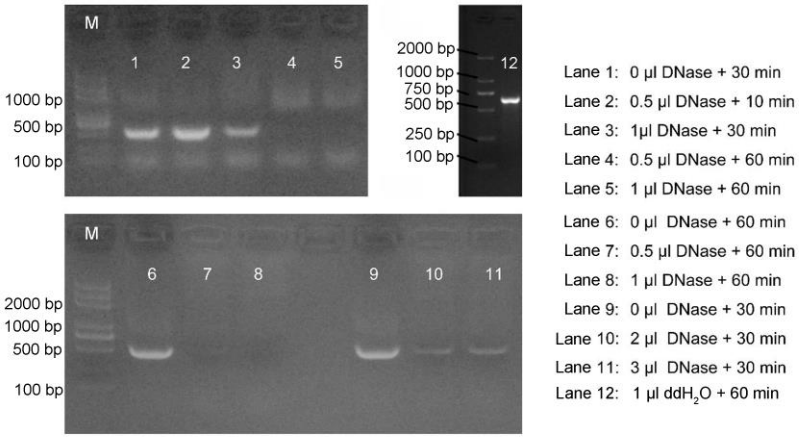

**Figure S3. Chilling treatments of two *Colpoda* species induced unstable cyst formation.** The chilling treatments at 0 ℃ and 4 ℃ showed that cold shock could induce formation of unstable cysts in actively growing *Colpoda* cells. Nevertheless, this effect was strongly dependent on chilling temperature and duration. Putting both *Colpoda* species at 10 ℃ did not induce formation of unstable cysts (data not shown). **(A)** Cell division of *Colpoda steinii* was completely inhibited upon ice water chilling, and nearly 100% of vegetative cells were transformed into unstable cysts in 4 hrs. Longer chilling duration led to a decrease in the number of unstable cysts. **(B)** In contrast, the population of *C. inflata* collapsed completely during ice water treatments within 3 hrs. **(C)** At 4 ℃, the small-sized *C. steinii* continued to grow and divide during the first 1.5 hrs. Afterwards, total cell abundance (= vegetative cells + unstable cysts) remained relatively stable, with approximately 76% of cells being transformed from vegetative status into unstable cysts following incubation for 9 to 12 hrs. Treating cells at 4 ℃ for a longer period of time damaged both types of cell. **(D)** Chilling at 4 ℃ completely inhibited cell division of *C. inflata*; unstable cysts progressively increased to 37% in 10 hrs, after which the number of unstable cysts remained stable for the next 8 hrs. All individuals transformed into unstable cysts in 20 hrs. Approximately, 36% of cells were lost during this incubation.

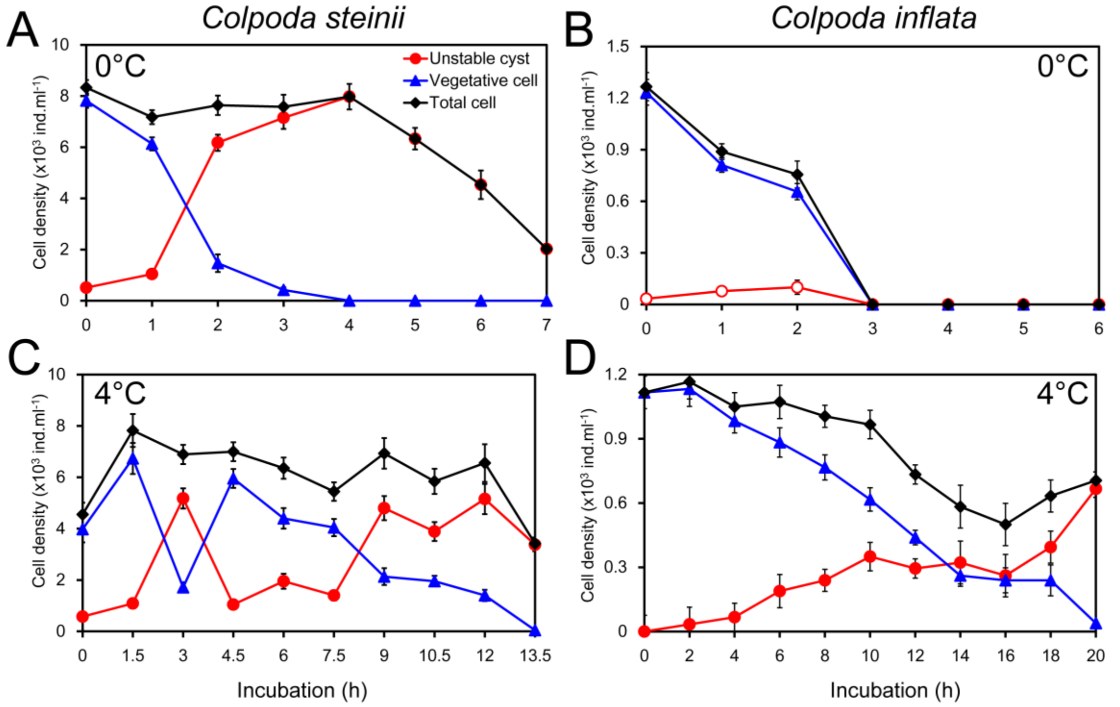

**Figure S4.** The per-cell rDNA and rRNA copy numbers (CNs) in two *Colpoda* species across life-cycle stages (lag, log, plateau phases and resting cysts) was significantly related to macronuclear volume. The rRNA CNs in RC appeared to be much lower than similarly sized vegetative cells or unstable cysts, such that the coefficients of determination ($R^2$) of regressions were much lower with resting cysts being taken into account (+RC) than without (-RC). RC = resting cyst.

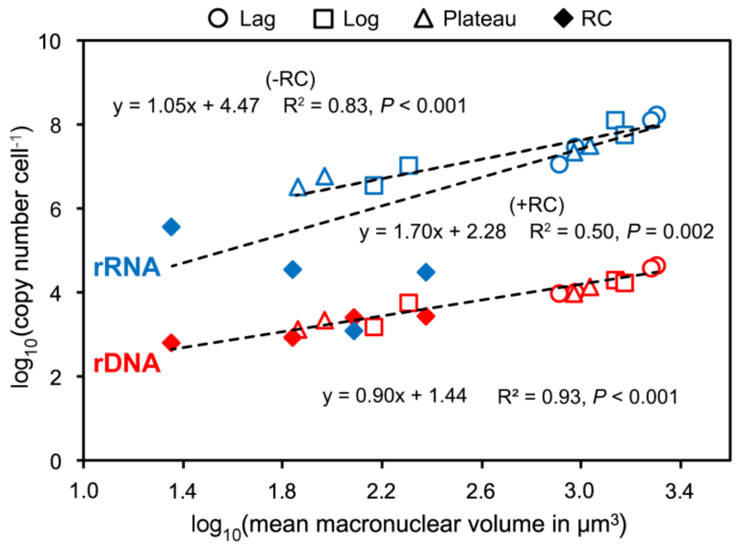

**Figure S5.** Regressions between rDNA and rRNA-based OTU numbers and cell volume.

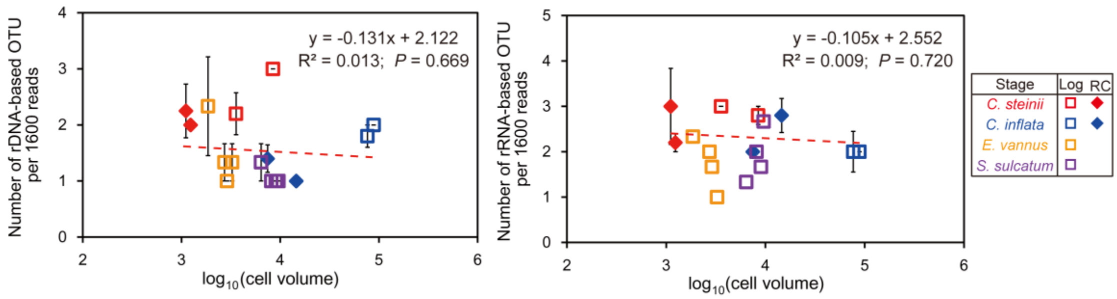

**Table S1. Numbers of reads (mean ± standard error) obtained at processing steps and resulting amplicon sequence variants (ASVs).**

| Species name | Life stage | Ribotype | Temp (℃) | Raw seqs | Filtered seqs | Merged seqs | Nonchimeras seqs | Retained reads (%)* | Range of OTU number# (All reads-based) | OTU number# (All reads-based) | OTU number# (Rarefied at 1600 reads) | *n* |
|---|---|---|---|---|---|---|---|---|---|---|---|---|
| *Colpoda steinii* | RC | DNA | 18 | 91999 ± 5186 | 78718 ± 3759 | 78160 ± 3689 | 77837 ± 3616 | 85 ± 2 | 2 ~ 3 | 2.3 ± 0.3 | 2.0 ± 0.0 | 3 |
| | | | 28 | 96381 ± 1323 | 85225 ± 2575 | 84805 ± 2674 | 84599 ± 2767 | 88 ± 2 | 1 ~ 4 | 2.5 ± 0.6 | 2.2 ± 0.1 | 4 |
| | | RNA | 18 | 89501 ± 3104 | 77448 ± 3135 | 76601 ± 3235 | 75742 ± 3505 | 85 ± 2 | 2 ~ 5 | 3.4 ± 0.5 | 2.0 ± 0.0 | 5 |
| | | | 28 | 91410 ± 2446 | 80311 ± 2601 | 79006 ± 2836 | 78873 ± 2833 | 86 ± 1 | 2 ~ 5 | 3.8 ± 0.6 | 2.9 ± 0.1 | 5 |
| | Log | DNA | 18 | 92836 ± 2137 | 81968 ± 1847 | 81630 ± 1843 | 81457 ± 1831 | 88 ± 0 | 3 ~ 3 | 3.0 ± 0.0 | 2.1 ± 0.0 | 5 |
| | | | 28 | 91359 ± 3405 | 80880 ± 3116 | 80600 ± 3130 | 80420 ± 3119 | 88 ± 1 | 1 ~ 4 | 2.8 ± 0.5 | 2.0 ± 0.0 | 5 |
| | | RNA | 18 | 86413 ± 1777 | 76451 ± 1612 | 75981 ± 1608 | 75854 ± 1591 | 88 ± 0 | 3 ~ 4 | 3.2 ± 0.2 | 2.1 ± 0.0 | 5 |
| | | | 28 | 90017 ± 3066 | 79539 ± 2526 | 78948 ± 2540 | 78623 ± 2555 | 87 ± 0 | 3 ~ 4 | 3.2 ± 0.2 | 2.1 ± 0.0 | 5 |
| *Colpoda inflata* | RC | DNA | 18 | 89468 ± 2956 | 78149 ± 2729 | 77865 ± 2730 | 77625 ± 2758 | 87 ± 1 | 1 ~ 2 | 1.2 ± 0.2 | 1.0 ± 0.0 | 5 |
| | | | 28 | 87593 ± 2039 | 75883 ± 1826 | 75366 ± 1893 | 74732 ± 1825 | 85 ± 0 | 1 ~ 3 | 1.6 ± 0.4 | 1.3 ± 0.0 | 5 |
| | | RNA | 18 | 92588 ± 2103 | 81082 ± 2102 | 80174 ± 2058 | 78524 ± 2138 | 85 ± 1 | 2 ~ 4 | 3.6 ± 0.4 | 2.3 ± 0.1 | 5 |
| | | | 28 | 90218 ± 3277 | 78868 ± 2550 | 77950 ± 2530 | 77067 ± 2511 | 85 ± 0 | 2 ~ 4 | 3.0 ± 0.3 | 2.0 ± 0.0 | 5 |
| | Log | DNA | 18 | 91796 ± 3184 | 82520 ± 2623 | 82312 ± 2663 | 82175 ± 2638 | 90 ± 0 | 2 ~ 2 | 2.0 ± 0.0 | 1.1 ± 0.0 | 4 |
| | | | 28 | 87413 ± 2711 | 78497 ± 2547 | 78186 ± 2558 | 78125 ± 2567 | 89 ± 0 | 1 ~ 4 | 2.4 ± 0.5 | 1.0 ± 0.0 | 5 |
| | | RNA | 18 | 90949 ± 2536 | 82514 ± 2489 | 82155 ± 2476 | 82095 ± 2482 | 90 ± 0 | 2 ~ 4 | 2.8 ± 0.5 | 1.1 ± 0.0 | 5 |
| | | | 28 | 93911 ± 2019 | 84351 ± 1635 | 84003 ± 1646 | 83959 ± 1631 | 89 ± 0 | 3 ~ 4 | 3.4 ± 0.2 | 1.1 ± 0.0 | 5 |
| *Euplotes vannus* | Log | DNA | 16 | 43540 ± 593 | 39011 ± 1425 | 38959 ± 1444 | 38926 ± 1457 | 90 ± 5 | 2 ~ 2 | 2.0 ± 0.0 | 1.0 ± 0.0 | 3 |
| | | RNA | | 43962 ± 6948 | 39696 ± 7819 | 39655 ± 7810 | 39651 ± 7809 | 89 ± 5 | 2 ~ 4 | 3.0 ± 0.6 | 1.4 ± 0.1 | 3 |
| | | DNA | 21 | 40096 ± 2735 | 35967 ± 4133 | 35934 ± 4135 | 35919 ± 4137 | 89 ± 5 | 2 ~ 4 | 3.0 ± 0.6 | 1.0 ± 0.0 | 3 |
| | | RNA | | 42861 ± 7676 | 38305 ± 8531 | 38251 ± 8514 | 38251 ± 8514 | 88 ± 6 | 1 ~ 4 | 3.0 ± 1.0 | 1.3 ± 0.1 | 3 |

| | | | | | | | | | | | | |
|---|---|---|---|---|---|---|---|---|---|---|---|---|
| | | DNA | 25 | 46696 ± 1993 | 41700 ± 3110 | 41639 ± 3122 | 41628 ± 3122 | 89 ± 5 | 3 ~ 5 | 4.0 ± 0.6 | 1.3 ± 0.1 | 3 |
| | | RNA | | 49308 ± 3910 | 44410 ± 5968 | 44346 ± 5968 | 44345 ± 5969 | 89 ± 5 | 1 ~ 6 | 3.3 ± 1.5 | 1.3 ± 0.1 | 3 |
| | | DNA | 16* | 38691 ± 3621 | 34395 ± 2613 | 34361 ± 2604 | 34360 ± 2603 | 89 ± 5 | 2 ~ 3 | 2.7 ± 0.3 | 1.0 ± 0.0 | 3 |
| | | RNA | | 43468 ± 1238 | 38958 ± 1616 | 38906 ± 1624 | 38904 ± 1623 | 90 ± 4 | 3 ~ 4 | 3.7 ± 0.3 | 2.3 ± 0.1 | 3 |
| *Strombidiu* | Log | DNA | 16 | 40614 ± 4207 | 33479 ± 3249 | 33424 ± 3236 | 33316 ± 3127 | 82 ± 1 | 2 ~ 3 | 2.5 ± 0.5 | 1.0 ± 0.0 | 2 |
| *m sulcatum* | | RNA | | 46633 ± 3220 | 41632 ± 5145 | 41559 ± 5173 | 41559 ± 5173 | 88 ± 5 | 3 ~ 5 | 4.3 ± 0.7 | 2.0 ± 0.2 | 3 |
| | | DNA | 21 | 37720 ± 3699 | 32927 ± 3997 | 32873 ± 4021 | 32778 ± 4115 | 87 ± 6 | 2 ~ 5 | 3.0 ± 1.0 | 1.3 ± 0.1 | 3 |
| | | RNA | | 42470 ± 3289 | 37398 ± 3879 | 37322 ± 3873 | 37313 ± 3882 | 88 ± 5 | 3 ~ 5 | 4.0 ± 0.6 | 2.0 ± 0.1 | 3 |
| | | DNA | 25 | 42498 ± 1378 | 37617 ± 3237 | 37572 ± 3247 | 37568 ± 3248 | 88 ± 5 | 2 ~ 6 | 4.0 ± 1.2 | 2.2 ± 0.1 | 3 |
| | | RNA | | 44956 ± 5466 | 40001 ± 7350 | 39924 ± 7377 | 39917 ± 7379 | 88 ± 5 | 2 ~ 4 | 3.0 ± 0.6 | 2.3 ± 0.1 | 3 |
| | | DNA | 16* | 39090 ± 1679 | 34395 ± 1670 | 34334 ± 1670 | 34319 ± 1667 | 88 ± 4 | 2 ~ 4 | 3.0 ± 0.6 | 1.3 ± 0.1 | 3 |
| | | RNA | | 44310 ± 3017 | 38509 ± 3954 | 38445 ± 3935 | 38445 ± 3935 | 86 ± 3 | 2 ~ 3 | 2.3 ± 0.3 | 1.0 ± 0.0 | 3 |

* Retained reads (%) indicate the proportion of retained reads with no chimeras to the number of raw reads of a sample.

[#], the OTUs are defined at a sequence similarity cutoff of 99% for *C. steinii* and *C. inflata*, and at 100% for *E. vannus* and *S. sulcatum*, based on the results of sequencing and analysis of individual clones with 18S rDNA or cDNA fragments.

*n*, the number of biological replicates.

**Table S2.** Performance of the DADA2 workflow in processing throughput sequencing data of randomly selected five individual clones inserted with 18S rDNA or cDNA (rRNA) fragments of four ciliate species. The numbers of unique amplicon sequence variants (ASVs, the reads that are 100% identical) and operational taxonomic units (OTUs) defined at a series of similarity threshold ranging from 99% to 90% are shown.

| **Species** | **Sample ID** | **ASV** | **OTU** | | | | | | | | | |
|---|---|---|---|---|---|---|---|---|---|---|---|---|
| | | **100** | **99** | **98** | **97** | **96** | **95** | **94** | **93** | **92** | **91** | **90** |
| *C. inflata* | CI1 | 12 | 1 | 1 | 1 | 1 | 1 | 1 | 1 | 1 | 1 | 1 |
| | CI2 | 12 | 1 | 1 | 1 | 1 | 1 | 1 | 1 | 1 | 1 | 1 |
| | CI3 | 12 | 1 | 1 | 1 | 1 | 1 | 1 | 1 | 1 | 1 | 1 |
| | CI4 | 12 | 1 | 1 | 1 | 1 | 1 | 1 | 1 | 1 | 1 | 1 |
| | CI5 | 12 | 1 | 1 | 1 | 1 | 1 | 1 | 1 | 1 | 1 | 1 |
| *C. steinii* | CS1 | 12 | 1 | 1 | 1 | 1 | 1 | 1 | 1 | 1 | 1 | 1 |
| | CS2 | 12 | 1 | 1 | 1 | 1 | 1 | 1 | 1 | 1 | 1 | 1 |
| | CS3 | 12 | 1 | 1 | 1 | 1 | 1 | 1 | 1 | 1 | 1 | 1 |
| | CS4 | 12 | 1 | 1 | 1 | 1 | 1 | 1 | 1 | 1 | 1 | 1 |
| | CS5 | 12 | 1 | 1 | 1 | 1 | 1 | 1 | 1 | 1 | 1 | 1 |
| *E. vannus* | EV1 | 1 | 1 | 1 | 1 | 1 | 1 | 1 | 1 | 1 | 1 | 1 |
| | EV2 | 1 | 1 | 1 | 1 | 1 | 1 | 1 | 1 | 1 | 1 | 1 |
| | EV3 | 1 | 1 | 1 | 1 | 1 | 1 | 1 | 1 | 1 | 1 | 1 |
| | EV4 | 1 | 1 | 1 | 1 | 1 | 1 | 1 | 1 | 1 | 1 | 1 |
| | EV5 | 1 | 1 | 1 | 1 | 1 | 1 | 1 | 1 | 1 | 1 | 1 |
| *S. sulcatum* | SS1 | 1 | 1 | 1 | 1 | 1 | 1 | 1 | 1 | 1 | 1 | 1 |
| | SS2 | 1 | 1 | 1 | 1 | 1 | 1 | 1 | 1 | 1 | 1 | 1 |
| | SS3 | 1 | 1 | 1 | 1 | 1 | 1 | 1 | 1 | 1 | 1 | 1 |
| | SS4 | 1 | 1 | 1 | 1 | 1 | 1 | 1 | 1 | 1 | 1 | 1 |
| | SS5 | 1 | 1 | 1 | 1 | 1 | 1 | 1 | 1 | 1 | 1 | 1 |
